# Fusion-associated sexual development in a testate amoeba fills a major gap in the evolution of sex in Amoebozoa

**DOI:** 10.64898/2026.07.05.736552

**Authors:** Yonas I. Tekle

## Abstract

Sexual processes in microbial eukaryotes are often cryptic, obscuring the diversity and evolutionary history of sex across major eukaryotic lineages. Within Amoebozoa, trophic-cell fusion has been associated with sexual development in distantly related taxa, but evidence from Tubulinea, one of the three major amoebozoan lineages, has been lacking, leaving a major gap in the known distribution of fusion-associated sexual development. Here, we combine long-term behavioral observations with transcriptomic analyses to uncover an extensive fusion-associated developmental program in the testate amoeba *Arcella vulgaris*. Individual trophic cells progressively fused with neighboring amoebae to form large multinucleate aggregates exhibiting coordinated movement and cytoplasmic streaming. Transcriptomic analyses identified a distinct meiosis-enriched state characterized by elevated expression of conserved meiotic genes, including DMC1, HOP1, HOP2, MER3, MSH5, REC8, ZIP4, and PCH2, together with genes involved in homologous recombination and chromosome maintenance. Morphologically similar fused aggregates occurred in both meiosis-enriched and meiosis-reduced transcriptomic states, revealing substantial molecular differentiation within the fusion process and suggesting a dynamic developmental continuum. The coordinated activation of conserved meiotic pathways strongly supports a role for trophic-cell fusion in sexual development. By extending fusion-associated sexual development to Tubulinea, our findings fill a major phylogenetic gap and establish the occurrence of this developmental phenomenon across all three major amoebozoan lineages. This broad phylogenetic distribution raises the possibility that fusion-mediated sexual development is an ancient and widespread feature of Amoebozoa and provides new insight into the evolution and diversity of sexual programs in microbial eukaryotes.

## Introduction

Microbial eukaryotes exhibit remarkable diversity in their reproductive and developmental strategies, yet sexual processes in many protistan lineages remain poorly characterized because they are often cryptic, episodic, or expressed only under specific environmental or developmental conditions ^1,2^. Over the past two decades, comparative genomic and transcriptomic studies have fundamentally challenged the long-standing assumption that many microbial eukaryotes are ancestrally asexual. Instead, surveys across diverse protistan groups have revealed widespread retention and expression of conserved meiotic machinery, indicating that sexual or parasexual processes are likely far more common than direct observations alone would suggest ^3–5^. Among microbial eukaryotes, Amoebozoa have emerged as one of the strongest examples of this pattern. Comparative analyses across the supergroup have demonstrated widespread conservation of genes associated with homologous recombination, chromosome pairing, and ploidy reduction, providing compelling evidence that sexual-developmental pathways are broadly retained throughout Amoebozoa ^1,6,7^. Consequently, the central question in amoebozoan biology has shifted from whether these organisms possess the genetic capacity for sex to when, how, and under what developmental contexts these pathways are activated. Despite this growing molecular evidence, however, direct stage-resolved observations linking cellular behavior to sexual-developmental programs remain available for only a limited number of amoebozoan lineages.

Among Amoebozoa, some of the strongest evidence for cryptic sexual development comes from taxa that undergo extensive cell fusion, multinucleate growth, or giant-cell formation. In the discosean amoeba *Cochliopodium*, repeated fusion among trophic amoebae produces large multinucleate plasmodial structures in which karyogamy, polyploid nuclei, and subsequent nuclear fragmentation have been documented using live-cell imaging and immunocytochemistry ^8^. Subsequent transcriptomic studies demonstrated that fused stages are enriched for meiosis- and karyogamy-associated genes, including DMC1, MER3, and PCH2, providing one of the clearest examples linking trophic cell fusion with sexual-developmental reprogramming in microbial eukaryotes ^9–11^. Comparable fusion-associated or giant-cell stages have also been reported in other amoebozoans, including *Entamoeba invadens*, where multinucleated giant cells arise during encystation and stress responses ^12^, dictyostelids in which mating-type-dependent cell fusion initiates macrocyst development ^13,14^, and myxogastrids where syngamy gives rise to multinucleate plasmodia ^15,16^. Together, these systems suggest that cell fusion, multinucleation, and giant-cell formation represent recurrent routes into sexual development across Amoebozoa rather than isolated lineage-specific phenomena. However, these observations remain phylogenetically fragmented, leaving unresolved whether fusion-associated sexual development is broadly conserved across the major amoebozoan divisions or represents multiple independent evolutionary innovations.

In contrast, sexual-developmental processes in Tubulinea remain comparatively obscure. Most tubulinean amoebae continue to be characterized primarily through morphology, ecology, and asexual binary fission, whereas direct evidence for sexual or fusion-associated developmental stages remains sparse and is derived largely from classical studies that have rarely been revisited using modern molecular approaches ^17,18^. This gap is particularly striking in Arcellinida (testate amoebae), a morphologically distinctive tubulinean lineage whose reproductive biology remains poorly understood despite decades of taxonomic, ecological, and evolutionary study ^1,19,20^.

Testate amoebae occupy a distinctive position in microbial eukaryote evolution because their preservable tests provide an unusually deep record of protistan morphological evolution ^21^. These tests contribute extensively to microfossil archives and paleoenvironmental reconstructions, making Arcellinida one of the few microbial eukaryote groups that provide a direct connection between extant protist biology and deep evolutionary history ^19,21^. Consequently, understanding life-cycle evolution in testate amoebae has implications that extend beyond contemporary microbial ecology to interpretations of the protistan fossil record and the evolutionary history of early eukaryotes. Despite their long history of taxonomic and ecological investigation, reproductive biology in testate amoebae remains poorly resolved. Classical studies of *Arcella* primarily focused on shell morphogenesis and binary fission, in which daughter cells construct new tests through highly coordinated secretion of Golgi-derived thecagenous granules before cytokinesis ^22–24^. Experimental studies further demonstrated that shell formation, cytokinesis, and body organization can become partially uncoupled during regeneration and exuviation-like processes, suggesting a greater degree of developmental plasticity than traditionally recognized ^25^. Direct evidence for sexuality in Arcellinida remains limited, but ultrastructural observations of reproductive cysts in *Arcella vulgaris* revealed synaptonemal complexes and meiotic-like nuclear reorganization associated with encystment ^17^. Additional reports of plasmogamy, karyogamy, and cyst-associated developmental transitions in *Arcella* and distantly related testate amoebae further suggest that sexual processes may occur more broadly within the group ^1,26^. However, these observations remain fragmentary, were largely made before the advent of modern molecular approaches, and have never been integrated with stage-resolved transcriptomic analyses capable of directly linking cellular behavior to activation of sexual-development pathways.

Here, we demonstrate that the testate amoeba *Arcella vulgaris* undergoes extensive trophic-stage cell fusion leading to the formation of large plasmodial-like aggregates involving numerous participating individuals. Comparative transcriptomic analyses reveal that these fusion stages are associated with activation of meiosis- and sex-related transcriptional programs, including genes involved in homologous recombination, chromosome pairing, meiotic progression, and chromosome maintenance. Together, these findings provide the first stage-resolved evidence of fusion-associated sexual development in Tubulinea, thereby closing a major phylogenetic gap in our understanding of sexual evolution across Amoebozoa. By extending this developmental program to the third major amoebozoan lineage, our results suggest that fusion-mediated sexual development may represent a deeply conserved feature of amoebozoan biology and provide a new framework for investigating the evolution of sexual life cycles in microbial eukaryotes.

## Materials and Methods

### *Arcella* Culture Conditions and Observation of Cellular Fusion

A monoclonal culture of *Arcella vulgaris* was established from a freshwater isolate and maintained in sterile Deer Park® spring water supplemented with varying numbers (1–10) of autoclaved rice grains to promote bacterial growth as a food source. Different rice grain densities were used to generate a range of nutrient conditions and population densities. Multiple independent cultures were maintained at room temperature and monitored over repeated approximately 30-day culture cycles for more than one year, allowing repeated observations of cellular behavior, aggregation, and fusion events under varying culture conditions.

Cellular behavior, aggregation, and fusion events were monitored using light microscopy following approaches previously employed in our studies ^8,9^. General morphology and cellular interactions were examined using an Eclipse Ti2 inverted microscope (Nikon Corporation, Japan). Still images and microscope-based time-lapse videos were acquired using the NIS-Elements software package under phase-contrast and differential interference contrast (DIC) optics. Cell measurements were obtained directly using NIS-Elements software. Additional long-term observations of culture behavior and fusion dynamics were documented using smartphone-based photography and videography.

Species identification was based on test morphology together with molecular data recovered from transcriptome assemblies, including small-subunit ribosomal RNA (SSU rRNA) and cytochrome oxidase subunit I (COI) sequences. Morphological characteristics and molecular data were consistent with previous descriptions of *Arcella vulgaris*.

Attempts were made to visualize nuclear behavior during fusion using DNA-staining approaches following Tekle *et al.* ^8^. However, reliable nuclear visualization was hindered by several technical challenges associated with *Arcella*. The shell (test) limited consistent penetration and distribution of fluorescent stains, while residual stain frequently accumulated within the test, resulting in high background fluorescence during imaging. In addition, the non-adherent and often fragile nature of fused aggregates complicated fixation and washing procedures required for fluorescence microscopy. Consequently, the present study focuses on cellular behavior and transcriptomic evidence rather than direct characterization of nuclear dynamics.

### RNA Isolation and Transcriptome Sequencing

Individual cellular samples representing distinct stages along the fusion continuum were isolated microscopically and transferred by mouth pipetting into sterile 0.2 mL PCR tubes for transcriptome sequencing. Samples included large fused aggregates, intermediate fusion stages, cells connected by cytoplasmic bridges, fully integrated plasmodial-like aggregates, and non-fused single-cell populations (Table S1). A total of twelve samples were collected for RNA sequencing. Two samples failed quality control and were excluded from downstream analyses, resulting in ten transcriptomes used for subsequent analyses (Table S1).

RNA extraction and cDNA synthesis were performed using the SMART-Seq v4 Ultra Low Input RNA Kit (Takara Bio USA) following the manufacturer’s protocol. cDNA concentrations were quantified using a Qubit Fluorometer and the Qubit DNA High Sensitivity Assay. Sequencing libraries were prepared by Azenta Life Sciences (Burlington, MA, USA) using the Nextera XT DNA Library Preparation Kit (Illumina Inc., San Diego, CA, USA) and sequenced on an Illumina platform using paired-end 150-bp chemistry (2 × 150 bp), generating approximately 10 million paired-end reads per sample.

Raw sequencing reads were assessed for sequence quality, read-length distribution, and adapter contamination using FastQC. Illumina adapter sequences and low-quality bases were removed using BBDuk, and reads failing quality filtering or becoming excessively short after trimming were discarded. The resulting quality-filtered reads were assembled de novo using rnaSPAdes under default parameters ^27^, following procedures previously applied in our amoebozoan transcriptomic studies ^9^. Individual transcriptome assemblies were generated to recover sample-specific transcripts, whereas a combined master assembly constructed from all samples was used for comprehensive gene inventory, functional annotation, and as the reference transcriptome for downstream differential gene expression analyses.

RNA-seq datasets have been deposited in the NCBI BioProject database under accession number XXXXXXX (pending).

### Identification of Sexual Development Genes

A curated set of 94 genes associated with meiosis, homologous recombination, DNA repair, chromosome cohesion, plasmogamy, karyogamy, and nuclear congression was compiled from previous studies of sexual development in Amoebozoa and other microbial eukaryotes ^6,11^. Candidate homologs were identified through sequence similarity searches against individual transcriptome assemblies and the combined master assembly using a custom gene-discovery pipeline ^6,11^. Recovered candidate sequences were subsequently validated using phylogenetic analyses. Protein sequences were aligned with representative homologs from diverse eukaryotic taxa using MAFFT v7 ^28^. Maximum-likelihood phylogenetic trees were reconstructed using IQ-TREE2 ^29^ with automatic model selection and ultrafast bootstrap support. Candidate genes were accepted as orthologs only when they grouped within the expected orthologous gene family in phylogenetic analyses. Predicted proteins were further examined using InterProScan v5 ^30^ to verify the presence of conserved domains consistent with their annotated identities. Presence–absence inventories were compiled for each individual transcriptome and the combined master assembly to assess the distribution of sexual-development genes across the fusion continuum.

### Identification of Transcriptomic States

Presence–absence inventories of sexual-development genes revealed substantial variation among transcriptomes in the recovery of meiosis-, recombination-, plasmogamy-, and karyogamy-associated genes. To determine whether this variation reflected distinct biological states, transcriptomes were further analyzed using principal component analysis (PCA) and hierarchical clustering of transcriptome-wide expression profiles.

PCA and hierarchical clustering consistently resolved two major transcriptomic groups. These groups were subsequently characterized according to the recovery and expression of meiosis-associated genes. One group exhibited broadly greater recovery and expression of meiosis-associated genes and was designated the Meiosis-Enriched State (**MES**), whereas the second exhibited comparatively reduced recovery and expression of these genes and was designated the Meiosis-Reduced State (**MRS**). Importantly, the two groups were resolved from transcriptome-wide expression structure and were not defined a priori using morphology or individual meiosis-associated genes.

### Transcriptomic Analyses

Differential gene expression (DGE) analyses were conducted to investigate transcriptional differences between the Meiosis-Enriched State (MES) and Meiosis-Reduced State (MRS). Transcript abundances were estimated by pseudoalignment of quality-filtered paired-end sequencing reads to the combined reference transcriptome using Kallisto v0.46 ^31^, with 100 bootstrap replicates generated for each sample. Transcript abundance estimates were imported into R using tximport ^32^ with txOut = TRUE and countsFromAbundance = “lengthScaledTPM”. Transcriptome-wide analyses were performed at the transcript level.

Differential expression analyses were performed using DESeq2 ^33^. Transcripts with fewer than 10 total estimated counts across all samples were excluded to minimize the influence of very low-abundance features. Size factors were estimated using the “poscounts” method implemented in DESeq2 to account for the sparse nature of transcript-level count data. Statistical significance was assessed using Wald tests, and P values were adjusted for multiple testing using the Benjamini–Hochberg false discovery rate (FDR) correction. Transcripts with an adjusted P value < 0.05 were considered significantly differentially expressed. Positive log₂ fold-change values indicate higher transcript abundance in the MES, whereas negative values indicate higher transcript abundance in the MRS.

Principal component analysis (PCA), hierarchical clustering, heatmap visualization, and permutational multivariate analysis of variance (PERMANOVA) were performed using variance-stabilized expression values generated by DESeq2 to assess transcriptome-wide expression structure and transcriptional variation among samples. PCA was used to summarize overall transcriptomic variation among samples, whereas hierarchical clustering was used to assess similarity in transcript expression profiles. Following identification and characterization of the two transcriptomic states, differentiation between MES and MRS was statistically evaluated using PERMANOVA implemented in the vegan package (adonis2) based on Euclidean distances with 999 permutations.

To visualize transcriptome-wide differential expression, MA plots were generated using ggplot2 by plotting the standard DESeq2 log₂ fold-change estimates against the log10-transformed mean normalized expression (*baseMean*) for each transcript. Meiosis-associated transcripts were highlighted and annotated according to their differential expression status, and labels were positioned using ggrepel to facilitate comparison with transcriptome-wide expression patterns.

Heatmaps were generated from variance-stabilized expression values using ComplexHeatmap, with color gradients produced using circlize::colorRamp2. Expression values were standardized by row-wise *z*-score normalization to visualize relative expression patterns among samples.

Genes were grouped according to their predicted biological functions, including meiosis, homologous recombination, DNA repair, chromosome cohesion, cell-cycle regulation, plasmogamy, karyogamy, and nuclear congression. Samples were ordered according to their transcriptomic state (MES and MRS).

### Comparative Analysis of *Arcella intermedia* Developmental Stages

To assess whether meiosis- and sexual-development-associated genes are expressed in other members of the genus, publicly available single-cell RNA-seq datasets from *Arcella intermedia* were reanalyzed. The dataset comprised single-cell transcriptomes from four growth phases (lag, log [exponential], stationary, and decline), with three single cells sampled from each growth phase, together with two whole-culture reference transcriptomes generated during construction of the *A. intermedia* reference transcriptome ^34^. The original study generated these transcriptomes to characterize transcriptional variation across the growth cycle and to construct a reference transcriptome for *A. intermedia*.

A curated set of meiosis- and sexual-development-associated genes identified in *Arcella vulgaris* and previous studies of sexual development in Amoebozoa was used as the reference gene set ^6^. Quality-filtered reads from each developmental phase and the two whole-culture reference transcriptomes were mapped directly to the reference gene set. Direct read mapping was employed because several target genes were not consistently recovered in de novo transcriptome assemblies, likely reflecting low transcript abundance and the stochastic nature of single-cell transcriptome assembly. Read-based analyses therefore provided a more sensitive assessment of target-gene expression across developmental stages.

Transcript quantification, normalization, differential expression analysis, and expression visualization followed the same transcript-level Kallisto, tximport, DESeq2, and variance-stabilizing transformation framework described above for *A. vulgaris*. Differential expression was evaluated across the lag, exponential, stationary, and decline growth phases, with transcripts having an adjusted *P* value < 0.05 considered significantly differentially expressed. The two whole-culture transcriptomes were included as references for visualization and comparative assessment but were not treated as developmental phases. Heatmaps were generated from variance-stabilized expression values standardized by row-wise z-score normalization, and significantly differentially expressed genes were indicated in the heatmap. The *A. intermedia* analysis was conducted independently of the *A. vulgaris* MES–MRS comparison and was used to assess temporal variation in sexual-development-associated gene expression across the growth cycle.

## Results

### Cellular Fusion and Formation of Large Plasmodia

Long-term observations of *Arcella vulgaris* cultures revealed frequent cell-cell interactions that resulted in progressive fusion among individuals (Figure 1; Supplementary Videos S1–S2). Fusion was initiated through direct contact between neighboring cells via cytoplasmic extensions emerging from the shell aperture. Following contact, plasma membrane fusion occurred rapidly, as evidenced by loss of the discernible boundary between interacting cells and the establishment of cytoplasmic continuity across the fusion site (Supplementary Video S1). Soon after fusion, cytoplasmic contents were observed to merge, forming broader regions of shared cytoplasm through which cytoplasmic streaming occurred while the connected cells continued to move and forage (Figure 1A–D, Supplementary Video S1).

**Figure 1.**
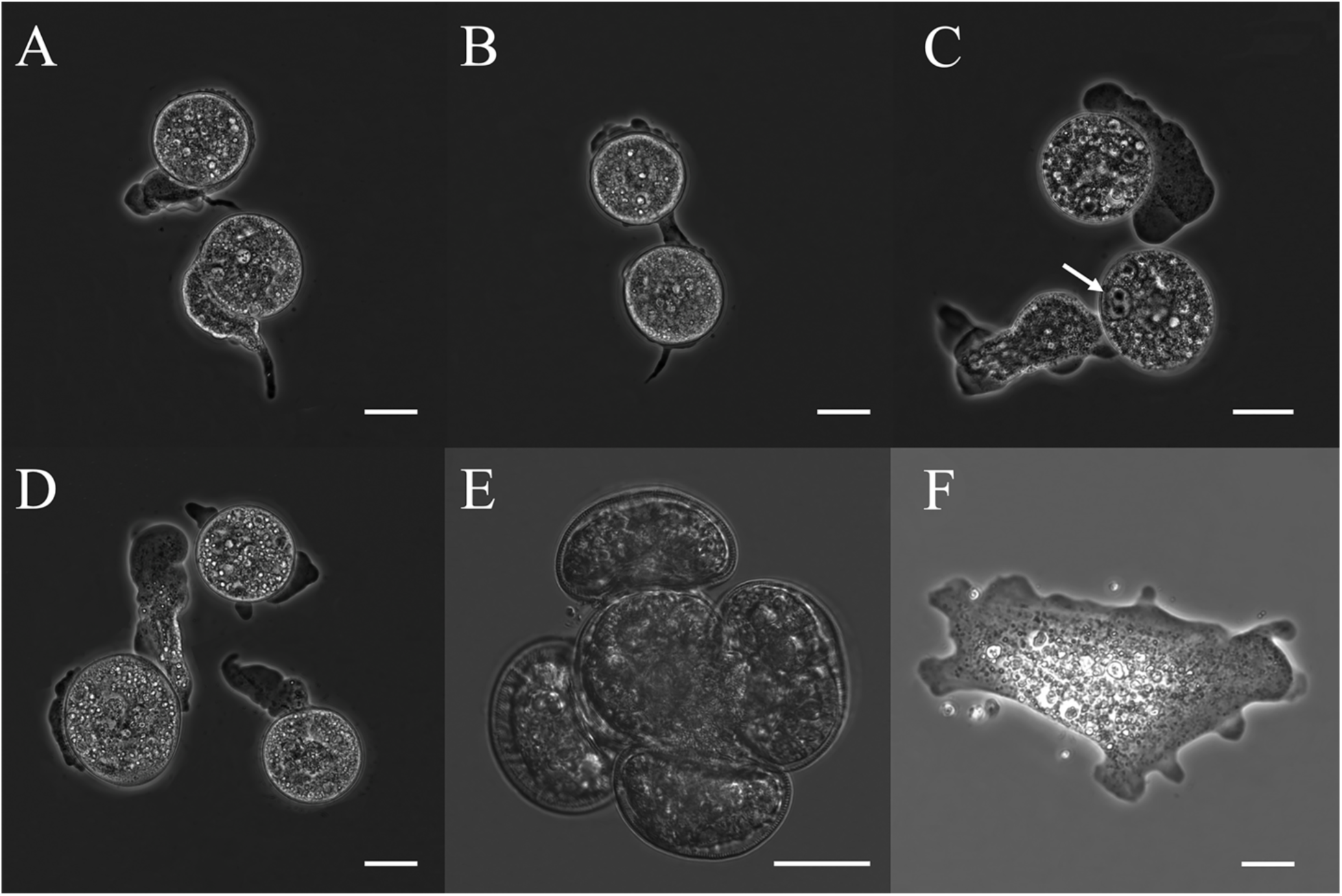
Early stages of trophic-cell fusion in *Arcella vulgaris*. (A–D) Representative micrographs showing initiation and progression of fusion between individual *Arcella* cells. The arrow in (C) indicates two closely associated nuclei within a likely fused cell, suggestive of a potential karyogamy-related stage. (E) Early fused aggregate formed through recruitment of multiple individuals. (F) Extruded multinucleated plasmodium lacking the test, displaying amoeboid morphology and extensive cytoplasmic integration. Scale bars = 20 μm.

Fusion events involved both pairwise interactions and larger multi-individual associations. Two-cell fusion events were common, but aggregates consisting of three or more individuals were also frequently observed (Figures 1, 2; Supplementary Videos S1–S2). Progressive incorporation of additional cells resulted in increasingly complex fused structures, indicating that fusion was not limited to isolated pairwise interactions but could proceed through repeated recruitment of neighboring individuals.

Advanced fusion stages were characterized by the formation of large plasmodia containing numerous attached shells distributed across the dorsal surface and periphery of the fused mass (Figure 2; Supplementary Videos S1–S2). Unlike individual trophozoites, these structures possessed a broad, unified cytoplasm exhibiting extensive coordinated streaming throughout the aggregate. Movement of the cytoplasmic mass often appeared eruptive or pulsatile, with large volumes of cytoplasm shifting across the fused structure in a manner not observed in unfused cells (Supplementary Video S2). Cells associated with shells in the dorsal region appeared integrated into a common cytoplasmic mass, whereas cells located along the periphery often retained recognizable pseudopodial morphology while remaining partially or fully fused to the aggregate. These peripheral cells frequently exhibited amoeboid activity and extended pseudopodia while maintaining cytoplasmic continuity with the larger fused structure. Collectively, the coordinated cytoplasmic streaming, persistent cytoplasmic continuity, and integrated movement patterns indicate that these structures behaved as single, integrated cellular entities rather than loose aggregations of interacting individuals.

**Figure 2.**
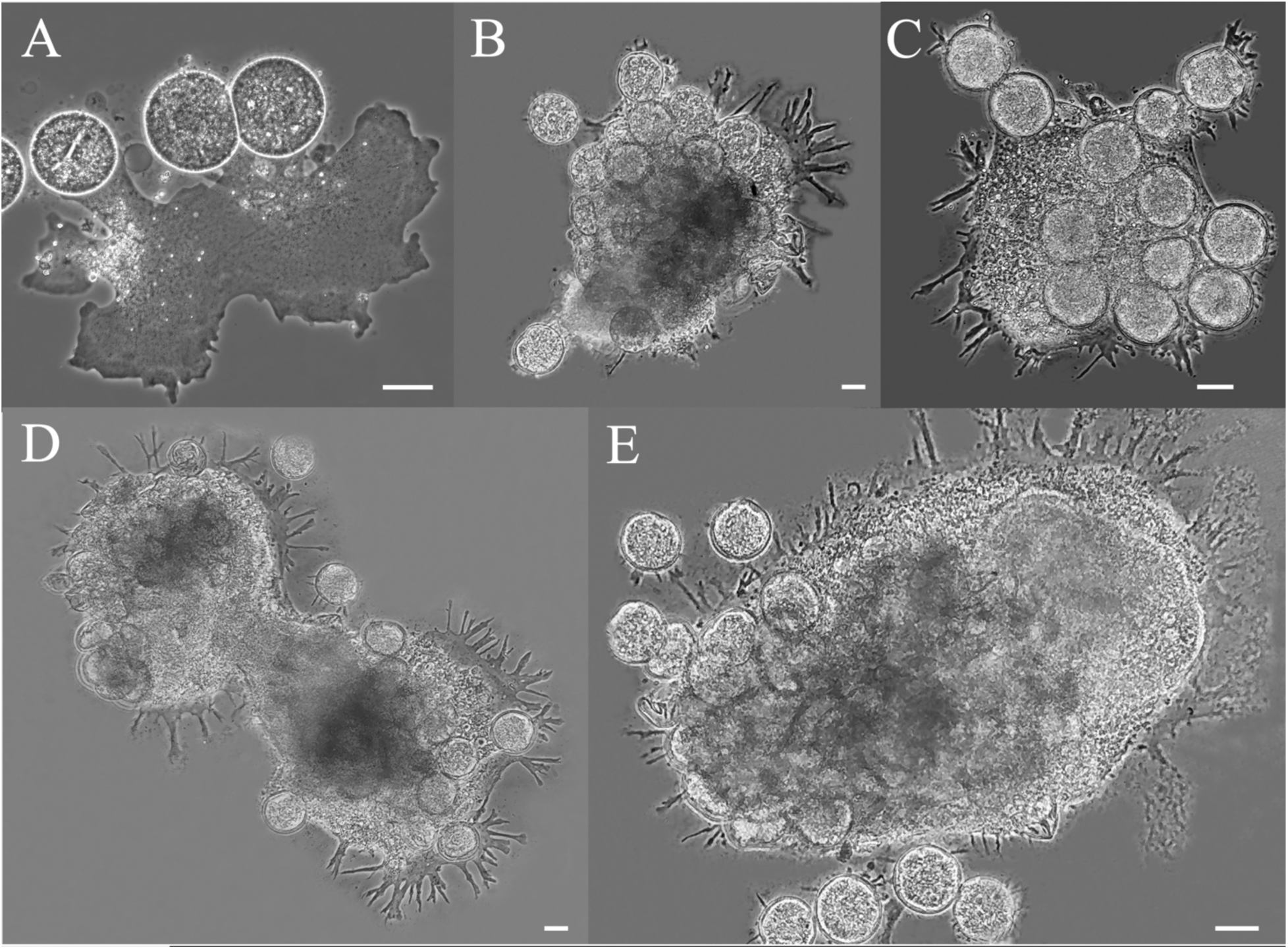
Formation of multinucleate giant-cell aggregates in *Arcella vulgaris*. Representative examples of advanced fusion stages resulting from progressive recruitment and integration of multiple individuals. (A) Early fused-cell stage with several attached cells sharing a common cytoplasm. (B–C) Intermediate fusion stages showing continued incorporation of additional cells and extensive peripheral pseudopodial activity. (D–E) Advanced multinucleate giant-cell aggregates containing numerous shells embedded within a shared cytoplasm. Scale bars = 20 μm.

Some aggregates contained more than 30 attached shells, indicating repeated fusion events involving numerous individuals (Figure 2A–E; Supplementary Video S2). In several instances, large plasmodia were observed merging with other plasmodia, demonstrating that fusion was not restricted to individual cells but could also occur between previously established fused structures (Supplementary Video S2). Large plasmodia remained viable and motile for periods ranging from approximately one to five days, during which they continued to forage, migrate, and recruit additional cells. Together, these observations demonstrate that cellular fusion in *A. vulgaris* is a progressive and dynamic process capable of generating large integrated cellular structures through repeated incorporation of multiple individuals and fusion among established plasmodia.

Individual trophozoites were commonly observed to contain up to three nuclei distributed throughout the cytoplasm. In some instances, two nuclei appeared in close proximity (Figure 1C). However, direct visualization of nuclear dynamics during and after fusion was limited by the test and dense, highly dynamic cytoplasm of large plasmodia, and attempts to monitor nuclear progression using live-cell staining and immunocytochemical approaches were unsuccessful (see Methods). Consequently, karyogamy and subsequent nuclear developmental stages could not be directly assessed in the present study.

### Inventory of Sexual Development Genes

Screening of transcriptome assemblies identified a substantial repertoire of genes associated with meiosis, homologous recombination, chromosome cohesion, DNA repair, plasmogamy, karyogamy, and nuclear congression (Table 1; Table S2). Overall, 11 of 13 surveyed meiosis-specific genes were recovered, including canonical components of the meiotic machinery such as DMC1, HOP1, HOP2, MER3/HFM1, MSH4, MSH5, PCH2, REC8, SPO11, and ZIP4. Phylogenetic analyses supported the orthology of recovered meiosis-, plasmogamy-, and karyogamy-associated genes (Figure S1).

**Table 1.** Summary of sexual-development genes detected in *Arcella vulgaris*.

| Functional Category | Genes Present | Representative Genes |
| --- | --- | --- |
| Meiosis-specific | 11/13 | DMC1, HOP1, HOP2, MER3, MSH4, MSH5, PCH2, REC8, SPO11, ZIP4 |
| Meiotic recombination and repair | 22/28 | RAD51, RAD54, MLH1, MLH3, PMS1, PMS2, MRE11, MUS81 |
| DNA damage response | 7/9 | MEC1/ATR, TEL1/ATM, RAD17, RAD24 |
| Sister chromatid cohesion | 7/8 | REC8, RAD21, SMC1–4 |
| Karyogamy | 3/6 | KAR2, JEM1, SEC63 |
| Plasmogamy-associated | 3/10 | MYO2, KEX2, KEM1 |
| Nuclear congression | 7/10 | CDC28, CDC34, KAR4, CIN4 |
| Cell polarization/congression | 2/2 | BNI1, TPM1 |

In addition to meiosis-specific genes, 22 of 28 genes associated with meiotic recombination and DNA repair pathways were identified, including RAD51, RAD54, MLH1, MLH3, PMS1, PMS2, MRE11, and MUS81. Genes associated with chromosome cohesion (7/8), DNA damage response (7/9), nuclear congression (7/10), karyogamy (3/6), and plasmogamy (3/10) were also detected (Table 1; Table S2).

Although many sexual-development genes were recovered from the combined master assembly, their distribution varied substantially among individual transcriptomes. Several meiosis-specific genes, including DMC1, REC8, SPO11, and ZIP4, were detected only in a subset of samples, whereas MER3, MSH5, PCH2, and the karyogamy-associated gene KAR2 were recovered from most transcriptomes (Table S2). This heterogeneous recovery of meiosis- and sexual-development-associated genes among samples suggested underlying transcriptional variation and prompted further analysis of transcriptomic differentiation among samples.

### Transcriptomic States and Differential Expression of Sexual Development Genes

Variation in gene inventories and transcriptome-wide expression profiles among samples suggested the presence of distinct biological states. Principal component analysis (PCA) based on transcriptome-wide expression profiles resolved two major transcriptomic groups (Figure 3). The first principal component explained 55% of the total variance, whereas the second explained 11%. These clusters corresponded to a Meiosis-Enriched State (MES) and a Meiosis-Reduced State (MRS). PERMANOVA analysis confirmed significant differentiation between the two states (R² = 0.287, P = 0.047).

**Figure 3.**
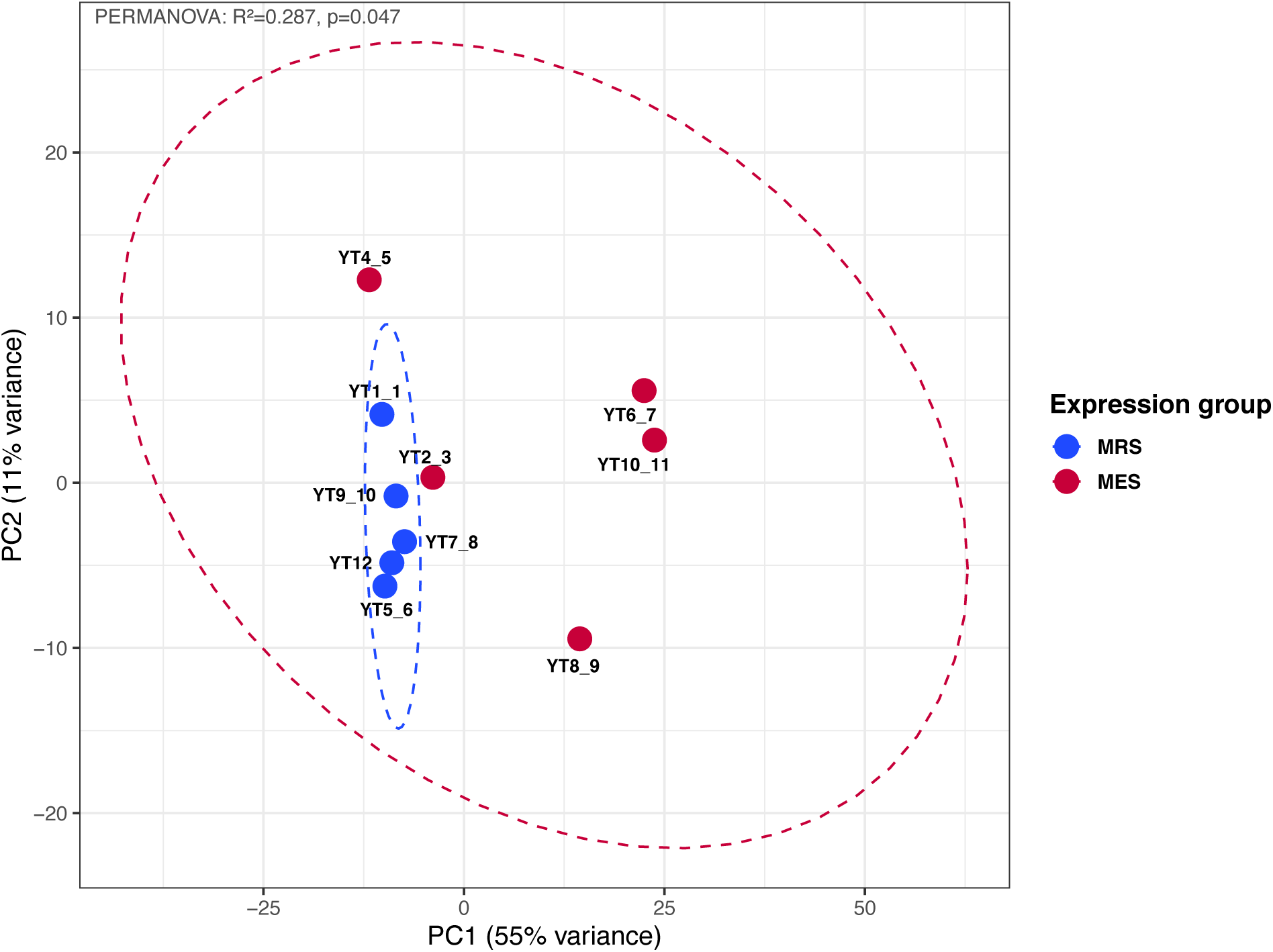
Principal component analysis (PCA) of transcriptome-wide expression profiles in *Arcella vulgaris*. Samples separate into two major groups corresponding to a Meiosis-Enriched State (MES) and a Meiosis-Reduced State (MRS). Morphologically similar fused samples occur in both groups, indicating that fusion encompasses multiple molecular states.

MES and MRS transcriptomes were not strictly associated with visible morphology. Samples with similar fused morphologies occurred in both transcriptomic states, indicating that morphological appearance alone does not fully capture molecular progression along the fusion continuum. Notably, both transcriptomic states contained fused samples, whereas the single unfused cell was assigned to the MRS, suggesting that substantial molecular differentiation occurs within the fusion continuum despite similar outward morphology.

Differential expression analyses further supported this distinction. The MA plot revealed an overall tendency toward higher expression of meiosis- and sex-related genes in MES transcriptomes (Figure 4). Many meiosis-associated genes, including DMC1, HOP1, MER3, MSH5, ZIP4, EXO1, RAD21, and RAD51, exhibited positive log₂ fold changes, although only a subset reached statistical significance. Additional genes involved in homologous recombination, DNA repair, chromosome cohesion, plasmogamy, and karyogamy likewise showed a general tendency toward elevated expression. Collectively, these patterns indicate a broad shift toward elevated expression of meiosis- and sexual-development-associated genes in MES transcriptomes rather than differential expression of only a few isolated genes.

**Figure 4.**
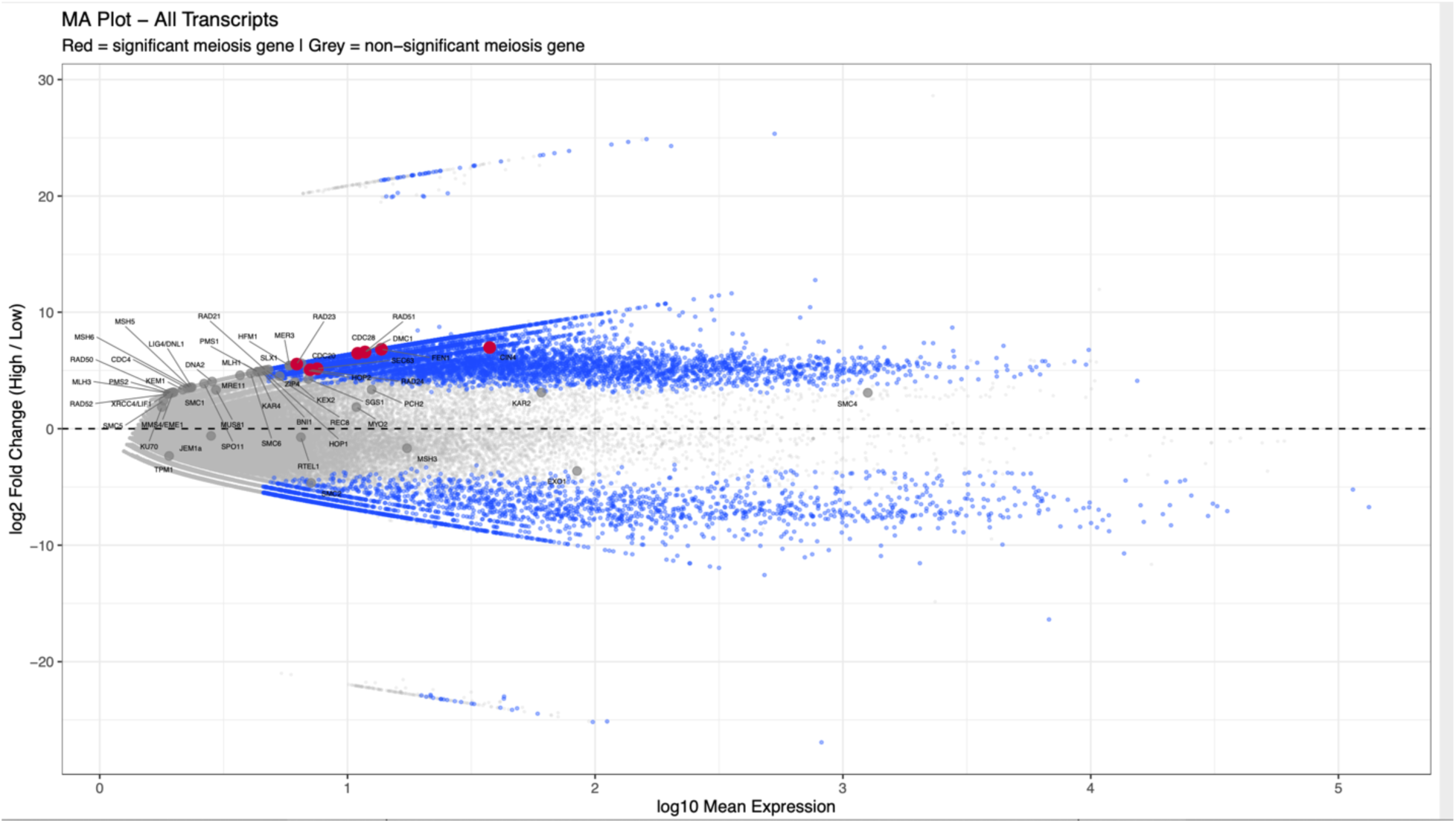
MA plot showing transcriptome-wide expression patterns, with meiosis- and sex-related genes highlighted. Each point represents a transcript plotted according to mean expression and log2 fold change. Red points indicate significantly differentially expressed genes, whereas gray points indicate non-significant meiosis- and sex-related genes. Most highlighted genes exhibit positive fold changes, consistent with overall enrichment of sexual-development pathways.

Heatmap analyses and hierarchical clustering revealed the same overall pattern, with MES and MRS transcriptomes forming distinct expression profiles (Figure 5; Figure S2). Meiosis-specific genes generally displayed lower expression in MRS transcriptomes and higher expression in MES transcriptomes. Particularly elevated expression in MES transcriptomes was observed for DMC1, MER3, HOP1, HOP2, MSH5, REC8, and ZIP4. Genes associated with plasmogamy and karyogamy also tended to exhibit higher expression in MES samples, including the plasmogamy-associated genes KEX2 and MYO2 and the karyogamy-associated genes JEM1, KAR2, and SEC63. Together, these analyses indicate a coordinated expression pattern involving genes associated with meiosis, homologous recombination, plasmogamy, karyogamy, and chromosome dynamics in MES transcriptomes.

**Figure 5.**
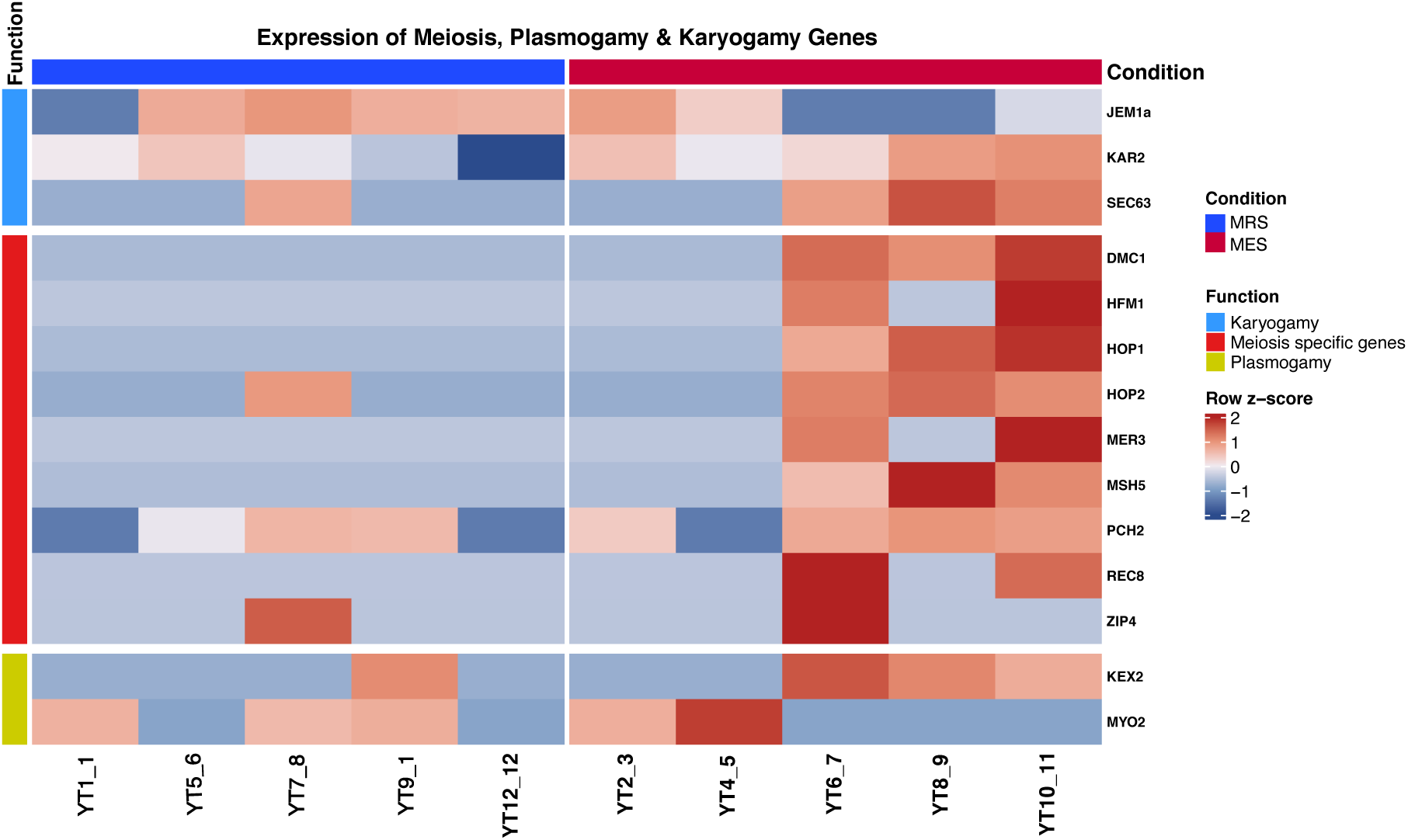
Heatmap showing normalized expression patterns of meiosis-, plasmogamy-, and karyogamy-associated genes in Meiosis-Enriched State (MES) and Meiosis-Reduced State (MRS) transcriptomes. Meiosis-specific genes, together with genes implicated in plasmogamy and karyogamy, exhibit coordinated enrichment in MES samples relative to MRS samples. Colors indicate row-standardized (z-score) expression values.

### Expression of Meiosis-Associated Genes During Growth of *Arcella intermedia*

Analysis of publicly available *Arcella intermedia* RNA-seq datasets revealed expression of multiple meiosis- and sexual-development-associated genes despite the absence of documented cellular fusion in this species (Figure S3). Direct read mapping detected expression of several meiosis-specific genes, including MER3, MND1, MSH5, HOP2, MSH4, and SPO11, together with genes associated with recombination, DNA repair, chromosome cohesion, plasmogamy, karyogamy, and nuclear congression.

Expression of many sexual-development-associated genes was elevated during the lag and log/exponential growth phases and generally reduced during the stationary and decline phases (Figure S3). The two whole-culture reference transcriptomes exhibited relatively uniform expression patterns across the genes examined, providing an additional reference for the expression of sexual-development-associated genes in mixed-stage cultures. Individual meiosis-and sexual-development-associated genes nevertheless showed variable expression across growth phases. Several genes were not recovered from transcriptome assemblies but were detected through direct read mapping, indicating that assembly-based inventories may underestimate low-abundance meiosis-associated transcripts. Together, these results suggest that expression of sexual-development-associated molecular pathways is elevated during active growth phases in *A. intermedia*.

## Discussion

### Fusion Defines a Sexual Developmental Program in *Arcella vulgaris*

The extensive trophic-cell fusion documented in *Arcella vulgaris* reveals a previously unrecognized developmental program in Tubulinea and provides the first direct evidence linking cellular fusion with molecular signatures of sexual development in a testate amoeba. Although fused *Arcella vulgaris* cells have previously been illustrated ^35^, the observation was not investigated further, and the biological or developmental significance of fusion remained unknown. Our observations demonstrate that fusion is neither an isolated interaction nor simple cellular aggregation. Individual trophozoites establish cytoplasmic continuity, progressively recruit additional cells, and form large, motile plasmodia containing a shared cytoplasmic mass. Some aggregates incorporated more than 30 individuals, and established plasmodia themselves fused with other plasmodia. Coordinated cytoplasmic streaming, persistent cytoplasmic continuity, and integrated movement further demonstrate that these structures behave as unified cellular entities (Figures 1, 2; Videos S1, S2).

The molecular data place this fusion behavior within a sexual developmental context. *A. vulgaris* possesses a broad repertoire of genes associated with meiosis, homologous recombination, chromosome cohesion, plasmogamy, karyogamy, and nuclear congression, including 11 of the 13 meiosis-specific genes surveyed. More importantly, these genes were not uniformly expressed across the fusion continuum. Transcriptome-wide analyses resolved two distinct molecular states, MES and MRS, with MES transcriptomes exhibiting broadly elevated expression of meiosis- and sexual-development-associated genes, including DMC1, HOP1, HOP2, MER3, MSH5, REC8, and ZIP4 (Figures 5, S2; Table S2). Although the presence or expression of meiosis-associated genes alone does not establish sexual development, their differential expression in the context of directly observed cellular fusion provides a fundamentally different level of evidence from gene inventories or constitutive expression alone ^3,4,36^. The convergence of cellular and transcriptomic evidence therefore supports the interpretation that trophic fusion in *A. vulgaris* is embedded within a sexual developmental program.

Notably, visible morphology did not reliably predict transcriptomic state. The unfused single-cell transcriptome was assigned to MRS (Figures 3, Table S1), whereas fused samples occurred in both MES and MRS. Thus, although the unfused state was associated with reduced meiosis-related expression, fusion morphology alone did not define the underlying molecular state. A similar disconnect between morphology and molecular state has been observed in *Cochliopodium*, where apparently comparable cells can occupy different pre-karyogamy, karyogamy, and post-karyogamy transcriptional states ^9^. MES and MRS may therefore represent molecularly distinct phases along a fusion-associated developmental trajectory rather than discrete morphological stages.

The nuclear events that follow fusion in *Arcella* remain unresolved. Direct visualization of nuclear behavior was limited by the test and the dense, highly dynamic cytoplasm of large plasmodia, preventing direct assessment of karyogamy or subsequent chromosome dynamics. Although molecular evidence cannot substitute for direct cytological observation of nuclear fusion, the coordinated expression of genes associated with nuclear congression, karyogamy, homologous recombination, and meiosis provides molecular support for the occurrence of additional nuclear developmental events following cytoplasmic fusion. Such events would be consistent with nuclear processes documented in the discosean amoeba *Cochliopodium* ^8,9^. Determining the sequence of nuclear interactions, ploidy transitions, and chromosome reorganization within *Arcella* plasmodia will be essential for resolving the complete developmental cycle. The present data, however, establish a direct association between extensive trophic-cell fusion and activation of a broad sexual-developmental molecular program in Tubulinea.

### Trophic Sexual Development Reveals an Overlooked Phase of the *Arcella* Life Cycle

The developmental timing of meiosis-associated gene expression provides an additional clue to the organization of the *Arcella* life cycle. The most influential evidence for sexual development in *Arcella* comes from classical ultrastructural studies of *A. vulgaris* reproductive cysts, in which synaptonemal complexes and meiotic-like nuclear reorganization were documented ^17^. This work established cyst-associated development as the primary documented context for meiosis in *Arcella* and has remained a central reference for sexuality in the genus. Our observations reveal a different and previously overlooked developmental context, placing elevated expression of meiosis- and sexual-development-associated genes within a continuum of actively motile, feeding, and fusing trophic cells. These findings suggest that activation of sexual-development machinery begins before, or independently of, the cyst-associated stages described in classical studies.

Independent transcriptomic data from *Arcella intermedia*, originally generated to characterize transcriptional variation across lag, exponential, stationary, and decline growth phases ^34^, provide an additional comparative context for this interpretation. Our analysis of these data showed that genes associated with meiosis, homologous recombination, chromosome cohesion, plasmogamy, karyogamy, and nuclear congression were expressed across the *A. intermedia* growth cycle. Many exhibited elevated expression during lag and exponential growth and generally reduced expression during stationary and decline phases (Figure S3). Thus, as in *A. vulgaris*, activation of sexual-development-associated machinery appears to coincide with active trophic growth rather than being restricted to late-stage culture development or conditions typically associated with encystment. The two whole-culture reference transcriptomes further showed broad expression across the surveyed sexual-developmental functional categories, indicating that these molecular programs are robustly represented within mixed *A. intermedia* cultures. These patterns broaden the developmental context of meiotic activity beyond the reproductive cyst stages documented in classical studies and are consistent with a trophic component of sexual-developmental activity.

The broad detection of sexual-development machinery in whole-culture *A. intermedia* transcriptomes also raises the possibility that fusion-competent or fusion-associated cells were present but unrecognized (Figure S3, Table S2). Fusion in *A. vulgaris* is transient, heterogeneous in appearance, and not reliably predicted by outward morphology, suggesting that comparable events could be overlooked in cultures examined primarily to characterize population growth and transcriptional variation. Alternatively, *A. intermedia* may activate a related sexual-developmental program without undergoing the extensive trophic-cell fusion observed in *A. vulgaris*. Distinguishing between these possibilities will require direct, long-term observation of *A. intermedia* cultures coupled with targeted sampling of cells expressing meiosis-associated genes.

Together, the *A. vulgaris* and *A. intermedia* data suggest that sexual development in *Arcella* may extend across a broader portion of the trophic life cycle than previously recognized. Trophic fusion could represent an early phase that precedes cyst-associated meiotic development, or fusion and encystment may represent alternative developmental routes into sexual processes. Although the relationship between trophic fusion and reproductive cyst formation remains unresolved, our findings substantially expand the developmental framework established by the classical cyst studies and identify actively growing trophic populations as an overlooked context for sexual development in *Arcella*.

### Fusion as a Deeply Rooted Route into Sexual Development Across Amoebozoa

The evolutionary significance of fusion-associated sexual development in *Arcella* becomes particularly apparent within the phylogenetic framework of Amoebozoa. Comparative genomic studies have demonstrated widespread retention of meiotic machinery across Amoebozoa and support the conclusion that the amoebozoan ancestor possessed the capacity for sexual reproduction, consistent with the deep ancestral origin of sex inferred across eukaryotes ^1,4,6,7,37^. Yet direct observations linking cellular behavior to sexual-developmental molecular programs have remained phylogenetically uneven. Discosea and Evosea contain several well-characterized fusion-associated sexual systems, whereas comparable modern evidence has been lacking from Tubulinea.

An intriguing historical exception is *Trichosphaerium*, an enigmatic amoebozoan lineage whose phylogenetic position remained uncertain for much of its taxonomic history and was only recently placed within Tubulinea by phylogenomic analyses ^38,39^. Classical studies proposed an alternating life cycle involving gamete formation and fusion in *Trichosphaerium*, although this sexual cycle has never been independently confirmed ^40^. Recent omics analyses recovered a broad meiosis-associated gene repertoire and demonstrated expression of these genes in multinucleate life stages, providing molecular support for a complex but still unresolved life cycle ^41^. Thus, *Trichosphaerium* provides a provocative but unresolved indication of sexual development in a lineage now recognized as tubulinean. The discovery of extensive trophic fusion coupled with meiosis-enriched transcriptomic states in *Arcella* provides the first modern stage-resolved evidence linking directly observed cellular fusion with sexual-developmental molecular signatures in Tubulinea and substantially strengthens the evidence for fusion-associated sexual development across the major amoebozoan lineages. This finding is particularly notable in Arcellinida, a lineage with an extensive fossil record, because it reveals a complex sexual-developmental capacity in a group that provides an unusual window into the deep evolutionary history of Amoebozoa ^21,42^.

The closest parallel occurs in the discosean amoeba *Cochliopodium*. Repeated fusion among trophic cells produces multinucleate plasmodia in which karyogamy, polyploid nuclei, and subsequent nuclear fragmentation have been directly documented ^8^. Cytological and transcriptomic analyses further showed that fused stages span pre-karyogamy, karyogamy, and post-karyogamy phases and are associated with elevated expression of meiosis- and sex-related genes ^9^. The striking parallels between *Arcella* and *Cochliopodium*, including trophic-cell fusion, progressive recruitment of multiple individuals, plasmodial development, and differential expression of meiotic machinery, are particularly notable given the deep evolutionary separation between Tubulinea and Discosea ^42^.

Evosea contains additional variations on this developmental theme. In dictyostelids, mating-type-dependent cell fusion initiates zygote formation and macrocyst development ^13,14,43^. In myxogastrids, syngamy between amoeboflagellate cells produces a diploid zygote that develops into a multinucleate plasmodium ^15,44^. *Entamoeba invadens* forms multinucleated giant cells through repeated cell fusion during encystation, accompanied by extensive nuclear reorganization and broader encystation-associated activation of recombination-related pathways ^12,45,46^. These systems differ substantially in morphology, developmental context, and life history, but repeatedly couple cellular fusion with multinucleate development, genome interaction, and sexual or recombination-associated processes. The comparative literature therefore supports a recurring association between whole-cell fusion, multinucleate or plasmodial stages, and sexual development across Amoebozoa.

Fusion-associated sexual development is now documented or strongly implicated in representatives of Discosea, Evosea, and Tubulinea (Figure 6). This phylogenetic distribution is consistent with the hypothesis that the capacity to couple cellular fusion with sexual development arose early in amoebozoan evolution. Under this interpretation, trophic fusion in *Arcella* and *Cochliopodium*, macrocyst development in dictyostelids, giant-cell formation in *Entamoeba*, and syngamy followed by plasmodial growth in myxogastrids represent lineage-specific manifestations of a deeply rooted developmental capacity. This does not require that their complete developmental cycles are homologous in every detail. Rather, an ancestral capacity linking cellular fusion with genome interaction and ploidy change may have provided a developmental framework that was subsequently modified within individual lineages. Fusion may therefore represent a conserved route into sexual development, while the downstream sequence of nuclear fusion, meiotic recombination, ploidy reduction, and developmental resolution diversified across Amoebozoa. Although multiple independent origins cannot presently be excluded, the occurrence of related developmental patterns across the deepest divisions of a supergroup already inferred to be ancestrally sexual provides a compelling framework for testing deep conservation against repeated convergence.

**Figure 6.**
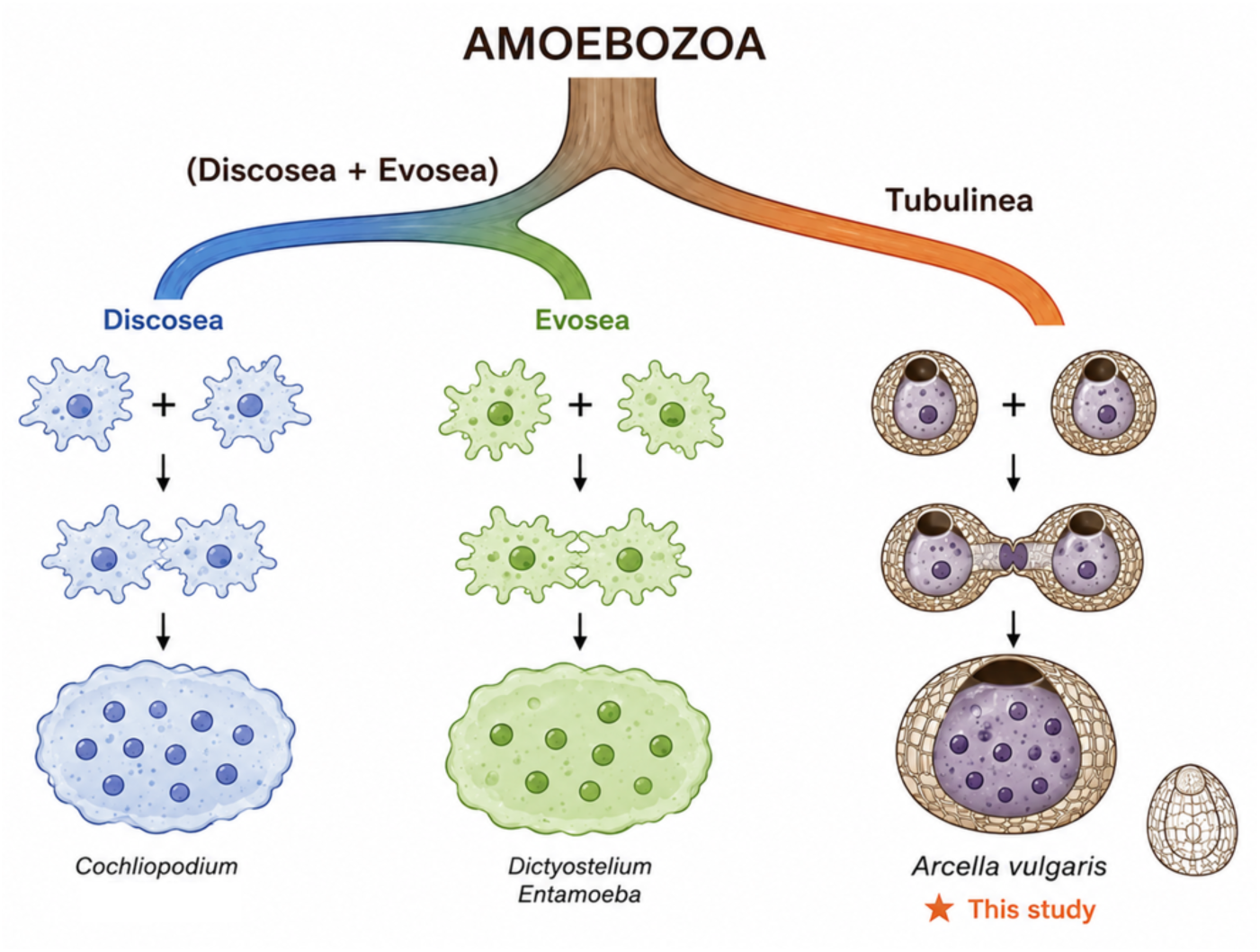
Fusion-mediated giant-cell formation across Amoebozoa. Representative examples of fusion and giant-cell formation in Discosea, Evosea, and Tubulinea. The discovery of fusion in *Arcella vulgaris* extends this phenomenon to all three major amoebozoan lineages.

The apparent rarity of fusion-associated development in many amoebozoans may also reflect observational bias. Sexual processes in microbial eukaryotes are often facultative, environmentally dependent, and difficult to capture through direct observation ^1,37^. In *Cochliopodium*, the occurrence and frequency of fusion vary among cultures and experimental observations ^8^, and our *Arcella* observations required repeated monitoring across culture cycles for more than one year. Long-term behavioral observation combined with stage-resolved transcriptomics may therefore reveal cryptic sexual-developmental processes in amoebozoan lineages currently regarded as lacking observable sex.

The apparent rarity of fusion-associated development in many amoebozoans may partly reflect observational bias. Sexual processes in microbial eukaryotes are often facultative, environmentally dependent, and difficult to capture through direct observation ^1,37^. In *Cochliopodium*, fusion behavior varies both among cultures and among members of the genus. Our observations across several species indicate substantial differences in the frequency and extent of fusion: some species fuse frequently and form large multinucleate plasmodia, whereas others fuse only rarely, despite retaining the capacity to do so ^8^. Similarly, recognition of the fusion-associated developmental behavior described here in *A. vulgaris* required repeated monitoring across culture cycles for more than one year. This extended observation period highlights how transient, infrequent, or taxon-specific variation in fusion behavior may remain undetected during shorter-term culture observations. Long-term behavioral observation combined with stage-resolved transcriptomics may therefore reveal cryptic sexual-developmental processes in amoebozoan lineages currently regarded as lacking observable sex.

### The Elusive Molecular Basis of Trophic-Cell Fusion

Despite the recurring association between cellular fusion and sexual development in Amoebozoa, the molecular machinery mediating trophic-cell fusion remains largely unknown. This gap is particularly striking in *Arcella* and *Cochliopodium*, where extensive fusion among actively growing trophic cells produces large integrated plasmodia. In both systems, fusion can involve repeated recruitment of multiple individuals and generate a shared cytoplasmic mass, yet the proteins responsible for cell recognition, membrane attachment, and plasma membrane fusion have not been identified ^7,10^. The conservation of such elaborate fusion behavior across deeply divergent amoebozoan lineages therefore contrasts sharply with our limited understanding of its molecular basis.

HAP2/GCS1 is an ancient and widely distributed eukaryotic gamete fusogen that mediates sexual cell fusion in diverse lineages and belongs to the fusexin superfamily of membrane fusion proteins ^47,48^. In Amoebozoa, HAP2/GCS1 homologs mediate gamete interactions in the dictyostelid *Dictyostelium discoideum* and have been detected in representatives of the three major amoebozoan lineages, although their distribution among sampled taxa is uneven ^7,47^. Intriguingly, HAP2/GCS1 is absent from the *Cochliopodium* genome ^10^, was not detected in the *Arcella intermedia* transcriptomic datasets examined here or previously surveyed by Hofstatter et al. (2018), and no convincing homolog was recovered from the *A. vulgaris* transcriptomes examined here, despite the extensive and repeated trophic-cell fusion observed in *Cochliopodium* and *A. vulgaris*. Although failure to recover HAP2/GCS1 from transcriptomic datasets cannot establish its genomic absence in *Arcella*, its repeated non-detection across independent transcriptomic datasets from two *Arcella* species, together with its absence from the *Cochliopodium* genome, raises the possibility that trophic-cell fusion in these amoebae may not depend on the canonical HAP2-mediated fusion pathway.

Such variation would not be unprecedented in amoebozoan sexual machinery. Individual canonical meiotic genes can be absent from amoebozoans with well-characterized sexual cycles; for example, DMC1 and SPO11 are absent from sampled dictyostelids despite their established sexual development ^6,7^. The absence or non-detection of HAP2/GCS1 therefore need not imply the absence of a sexual fusion process, but may instead reflect functional replacement, extreme sequence divergence, or recruitment of alternative membrane-fusion machinery. The molecular mechanism of plasmogamy may consequently be more evolutionarily labile than the developmental use of cellular fusion itself.

Fusion-associated multinucleate development may also extend beyond Amoebozoa. In a recently characterized heterolobosean isolate from Mombasa, Kenya, we observed cellular fusion, formation of multinucleate plasmodial-like cells containing polyploid nuclei, and subsequent fragmentation into smaller cellular units ^49^. Although the relationship of this developmental cycle to sex remains unresolved, its behavioral parallels with *Cochliopodium* and *Arcella* suggest that fusion followed by multinucleate or polyploid growth may be more broadly distributed among amoeboid microbial eukaryotes than currently recognized. The occurrence of similar cellular behaviors in deeply divergent lineages further raises the question of whether these organisms employ evolutionarily related fusion machinery or have independently recruited different molecular mechanisms to achieve comparable developmental transitions.

Identifying the proteins responsible for trophic-cell recognition and membrane fusion will be critical for distinguishing between these possibilities. If deeply divergent amoebozoan lineages use homologous fusion machinery, this would provide strong evidence for conservation of an ancestral molecular mechanism. Conversely, the use of unrelated fusogens would suggest repeated molecular innovation within a developmental framework in which cellular fusion remains a route into sexual development. Comparative analysis of fusion-enriched cellular states, particularly across *Arcella*, *Cochliopodium*, and other fusion-capable amoeboid lineages, may therefore provide a route to identifying previously unrecognized eukaryotic fusion machinery and resolving the evolutionary origin of trophic-cell plasmogamy. More broadly, the persistence of comparable fusion-associated developmental programs despite possible turnover in the underlying fusion machinery raises the possibility that the developmental role of cell fusion is more evolutionarily conserved than the specific proteins that execute membrane fusion. Together, our findings identify trophic-cell fusion as a previously overlooked component of sexual development in Tubulinea and support a broader model in which fusion represents a deeply rooted route into sexual development across Amoebozoa.

## Resource Availability

### Lead contact

Further information and requests for resources should be directed to and will be fulfilled by the lead contact, Yonas I. Tekle.

### Materials availability

This study did not generate new unique reagents. *Arcella vulgaris* cultures used in this study are available from the lead contact upon reasonable request.

### Data and code availability

RNA-sequencing data generated in this study have been deposited in the NCBI Sequence Read Archive (SRA) under BioProject accession number XXXXXXX and will be publicly available as of the date of publication. Publicly available *Arcella intermedia* RNA-sequencing datasets analyzed in this study were obtained from Ribeiro et al. (2020).

No original code was developed specifically for this study. Analyses were performed using publicly available software and previously described analytical workflows, as detailed in the Methods.

### Author Contributions

Y.I.T. conceived and designed the study, conducted the experiments and long-term behavioral observations, performed and supervised the transcriptomic and evolutionary analyses, interpreted the data, prepared the figures, and wrote and revised the manuscript.

## Acknowledgments

This work is supported by the National Institutes of Health (1R15GM116103-02), Simons Foundation Fellow Award (SFA-23-5) and National Science Foundation Excellence in Research (EiR) Award #2401946. The author thanks Priyal Patel for technical assistance with laboratory work and implementation of the transcriptomic analyses.

## Supplementary Figure captions

**Supplementary Figure S1.**
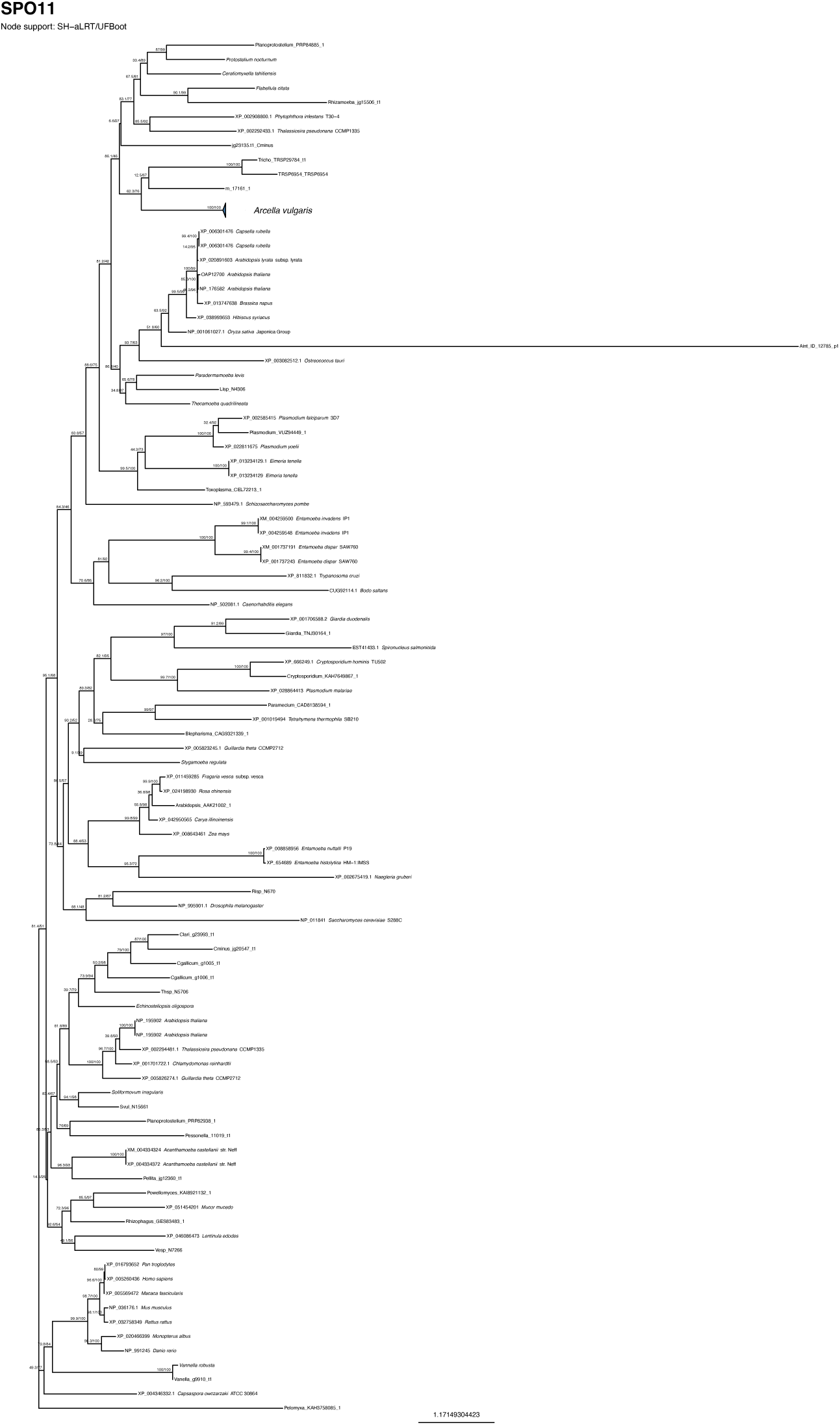
Phylogenetic validation of representative meiosis- and sex-related genes identified in *Arcella vulgaris*. Maximum-likelihood phylogenetic trees used to confirm the identity and orthology of representative meiosis- and sex-related genes recovered from *Arcella transcriptomes*. Branch support values represent ultrafast bootstrap percentages.

**Supplementary Figure S2.**
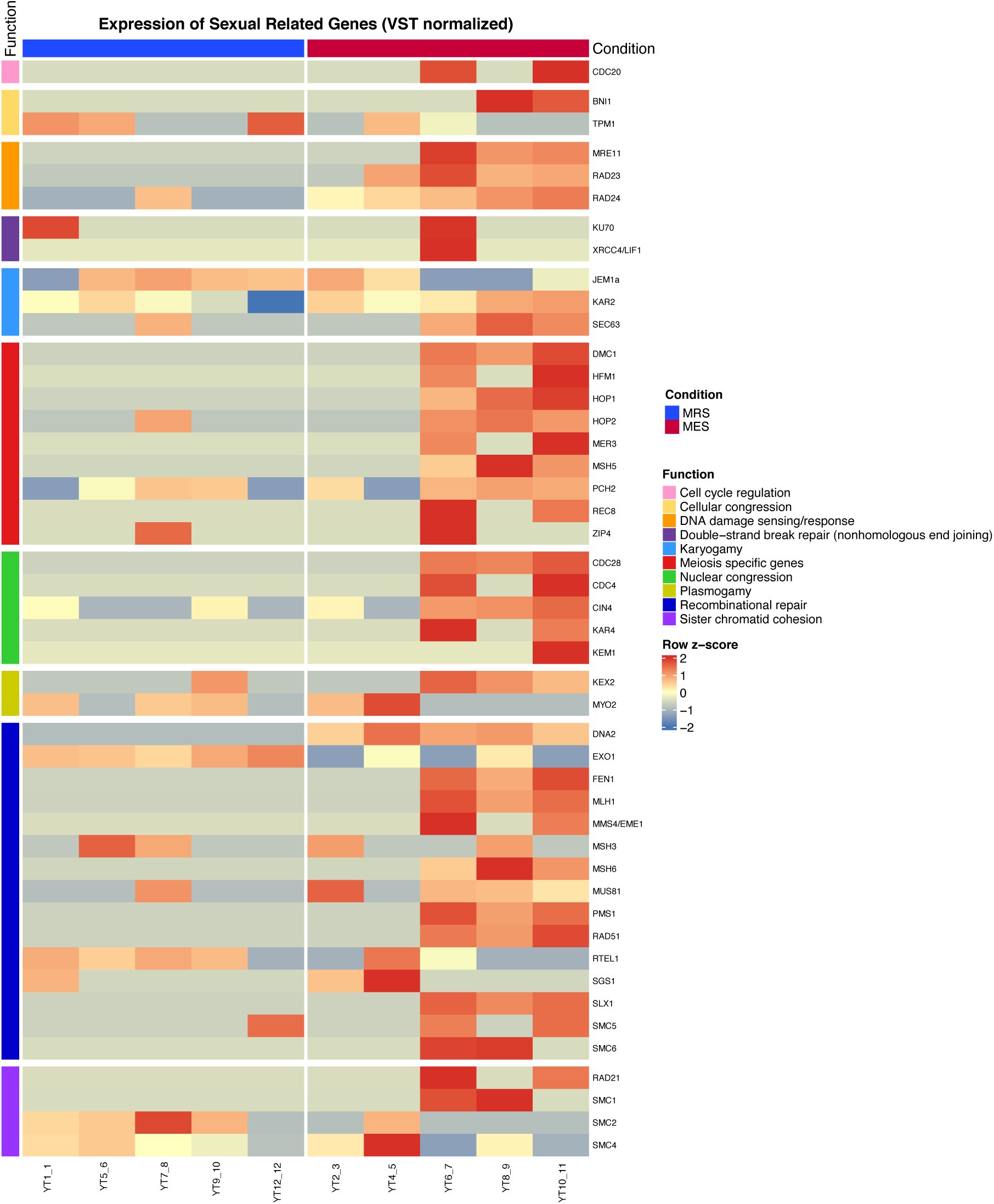
Expanded expression patterns of sexual-development genes across *Arcella vulgaris* transcriptomic states. Heatmap showing normalized expression of the complete set of sexual-development genes recovered from *Arcella vulgaris* transcriptomes. This figure expands upon Figure 5 by including genes associated with meiosis, recombination and DNA repair, plasmogamy, karyogamy, nuclear congression, sister chromatid cohesion, cell-cycle regulation, and DNA damage response pathways. Samples are grouped according to Meiosis-Enriched State (MES) and Meiosis-Reduced State (MRS) transcriptomic states. Colors indicate row-standardized (z-score) expression values.

**Supplementary Figure S3.**
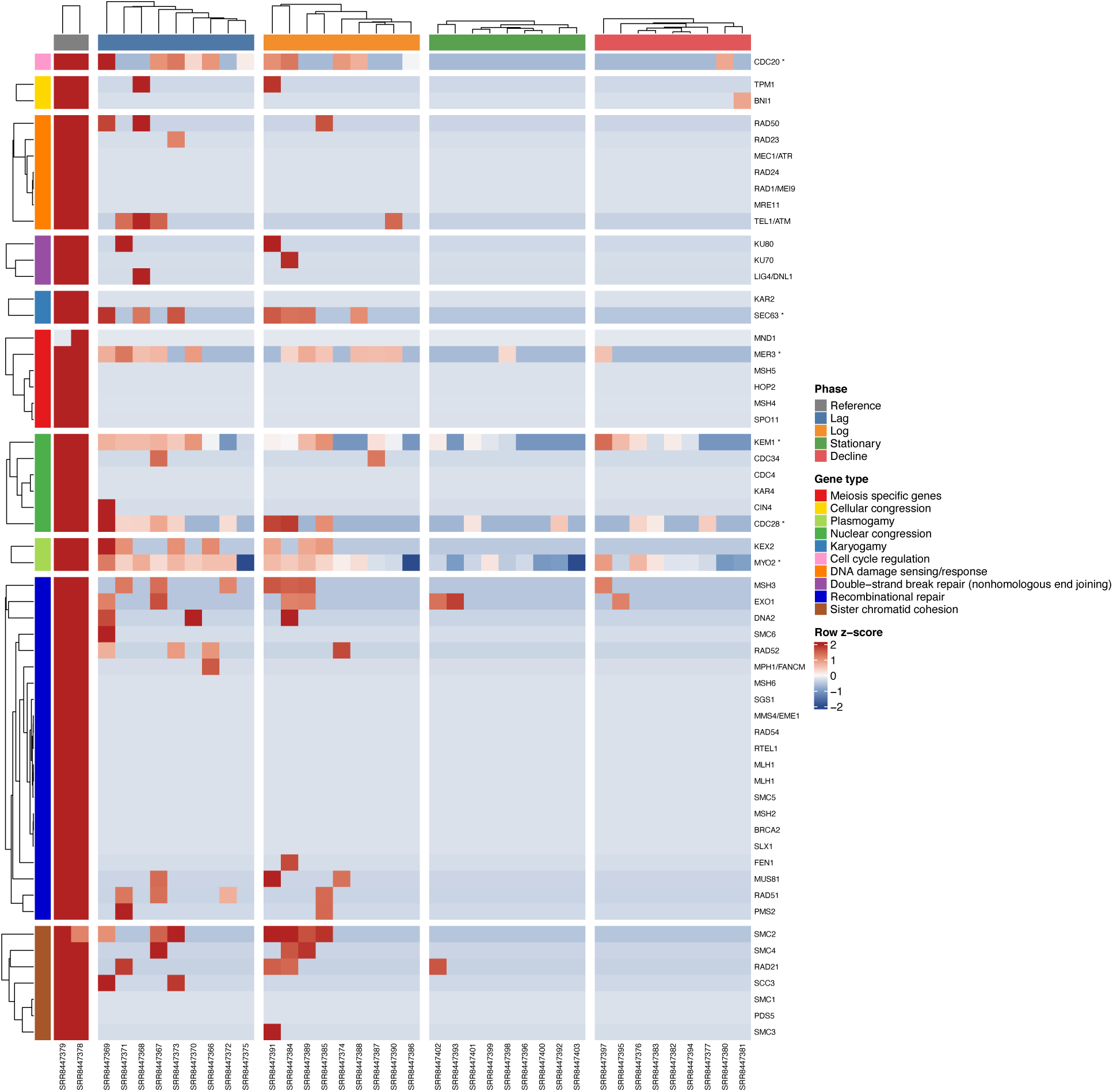
Expression of meiosis-associated genes across growth phases in *Arcella intermedia*. Heatmap showing expression patterns of meiosis- and sexual-development-associated genes across the Lag, Log (exponential), Stationary, and Decline growth phases, together with two whole-culture reference transcriptomes of *Arcella intermedia*. Genes identified as significantly differentially expressed are indicated by an asterisk (*). Most genes exhibit elevated expression during the Lag and Log phases and reduced expression during the Stationary and Decline phases, consistent with greater activation of meiosis- and sexual-development-associated pathways during active vegetative growth than during late-stage culture development. The whole-culture reference transcriptomes exhibit relatively uniform expression patterns across the genes examined. Colors represent row-standardized (z-score) expression values, and genes are grouped according to their predicted functional categories.

**Supplementary Video S1.** Initiation of trophic-cell fusion in *Arcella vulgaris*. Time-lapse video showing the initiation of fusion between individual *Arcella vulgaris* cells. Adjacent amoebae establish contact through their extended pseudopodia and progressively merge, resulting in cytoplasmic continuity between neighboring cells. This video corresponds to the early fusion stages illustrated in Figure 1 and documents the formation of the initial fused state. **YouTube link**: https://www.youtube.com/shorts/3lCNmmqPCa8

**Supplementary Video S2**. Coordinated movement and cytoplasmic integration in giant-cell aggregates of *Arcella vulgaris*. The collection of videos shows advanced fusion stages in various *Arcella vulgaris* plasmodia that are composed of multiple fused individuals sharing a common cytoplasm. Cells at the periphery join and leave the plasmodia while maintaining pseudopodial extensions, whereas numerous shells remain embedded across the dorsal surface of the dynamically streaming cytoplasmic mass. The videos illustrate the dynamic nature of the multinucleate giant-cell stages and their high degree of cellular integration achieved following multiple trophic-cell fusion. **YouTube link**: https://youtu.be/TnpcvzxKUM0

**Table S1.**
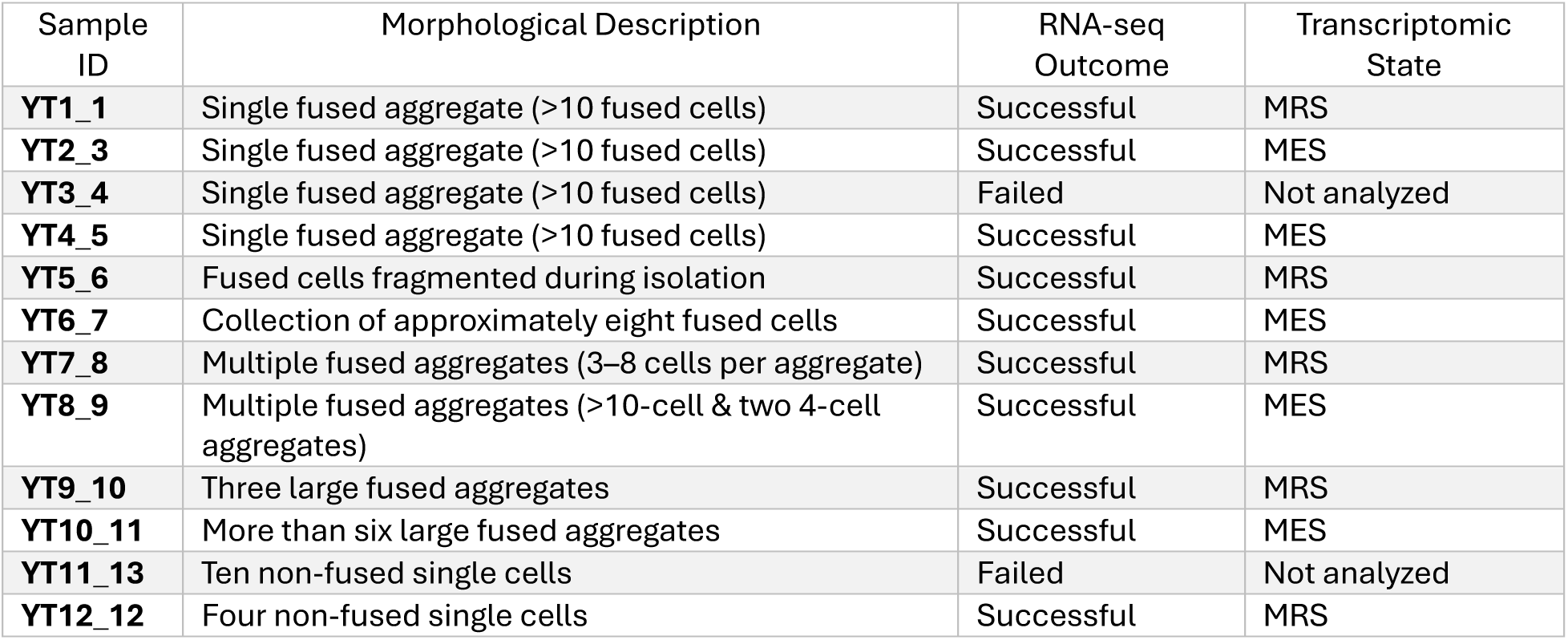
*Arcella vulgaris* transcriptomic samples used in this study and their assignment to transcriptomic states.

| Sample ID | Morphological Description | RNA-seq Outcome | Transcriptomic State |
| --- | --- | --- | --- |
| YT1_1 | Single fused aggregate (>10 fused cells) | Successful | MRS |
| YT2_3 | Single fused aggregate (>10 fused cells) | Successful | MES |
| YT3_4 | Single fused aggregate (>10 fused cells) | Failed | Not analyzed |
| YT4_5 | Single fused aggregate (>10 fused cells) | Successful | MES |
| YT5_6 | Fused cells fragmented during isolation | Successful | MRS |
| YT6_7 | Collection of approximately eight fused cells | Successful | MES |
| YT7_8 | Multiple fused aggregates (3–8 cells per aggregate) | Successful | MRS |
| YT8_9 | Multiple fused aggregates (>10-cell & two 4-cell aggregates) | Successful | MES |
| YT9_10 | Three large fused aggregates | Successful | MRS |
| YT10_11 | More than six large fused aggregates | Successful | MES |
| YT11_13 | Ten non-fused single cells | Failed | Not analyzed |
| YT12_12 | Four non-fused single cells | Successful | MRS |

**Supplementary Table S2.** *Inventory* of meiosis and sex related genes identified in *Arcella vulgaris* (MRS and MES) transcriptomes and whole culture based transcriptome of *Arcella intermedia* from ^34^ used here as a reference.

| Function | Gene | OG | *Arcella*<br>*intermedia*<br>*a* | Meiosis-Reduced State<br>(MRS) |  |  |  |  | Meiosis-Enriched State<br>(MES) |  |  |  |  |
| --- | --- | --- | --- | --- | --- | --- | --- | --- | --- | --- | --- | --- | --- |
|  |  |  |  | YT12<br>-12 | YT9-10 | YT1-1 | YT5-6 | YT7-8 | YT4-5 | YT2-3 | YT8-9 | YT6-7 | YT10<br>-11 |
| *Meiosis-specific* | *DMC1* | OG5_126834 | + | n<br>d | n<br>d | n<br>d | n<br>d | n<br>d | n<br>d | + | + | + | + |
|  | *MND1* | OG5_127882 | + | n<br>d | n<br>d | n<br>d | n<br>d | n<br>d | n<br>d | n<br>d | + | n<br>d | + |
|  | *HOP2* | OG5_128568 | + | n<br>d | n<br>d | + | + | n<br>d | + | + | + | + | + |
|  | *HOP1* | OG5_128667 | + | + | + | n<br>d | + | + | + | + | + | + | + |
|  | *MSH5* | OG5_129379 | + | + | + | + | + | + | + | + | + | + | + |
|  | *MER3* | OG5_129931 | + | + | + | n<br>d | + | n<br>d | + | n<br>d | + | + | + |
|  | *MSH4* | OG5_130077 | + | + | + | + | + | + | + | + | + | + | + |
|  | *ZIP-like* | OG5_171209 | nd | n<br>d | n<br>d | n<br>d | n<br>d | n<br>d | n<br>d | n<br>d | n<br>d | n<br>d | n<br>d |
|  | *RED1* | OG5_180525 | nd | n<br>d | n<br>d | n<br>d | n<br>d | n<br>d | n<br>d | n<br>d | n<br>d | n<br>d | n<br>d |
|  | *SPO11* | OG5_127274 | + | n<br>d | n<br>d | n<br>d | n<br>d | n<br>d | n<br>d | + | + | + | + |
|  | *REC8* | OG5_150817 | + | n<br>d | n<br>d | n<br>d | n<br>d | n<br>d | n<br>d | n<br>d | n<br>d | + | n<br>d |
|  | *PCH2* | OG5_128995 | + | + | + | + | + | + | + | + | + | + | + |

|  |  |  |  | Meiosis-Reduced State (MRS) |  |  |  |  | Meiosis-Enriched State (MES) |  |  |  |  |
| --- | --- | --- | --- | --- | --- | --- | --- | --- | --- | --- | --- | --- | --- |
| Function | Gene | OG | *Arcella intermedi*<br>*a* | YT12<br>-12 | YT9-10 | YT1-1 | YT5-6 | YT7-8 | YT4-5 | YT2-3 | YT8-9 | YT6-7 | YT10<br>-11 |
|  | *ZIP4* | OG5_142513 | + | n<br>d | n<br>d | n<br>d | n<br>d | n<br>d | n<br>d | + | + | + | + |
|  | *HAP2* | OG5_129105 | nd | n<br>d | n<br>d | n<br>d | n<br>d | n<br>d | n<br>d | n<br>d | n<br>d | n<br>d | n<br>d |
|  | *GEX1* | OG5_168665 | nd | n<br>d | n<br>d | n<br>d | n<br>d | n<br>d | n<br>d | n<br>d | n<br>d | n<br>d | n<br>d |
| Cellular<br>congression | *BNI1* | OG5_127406 | + | n<br>d | n<br>d | n<br>d | n<br>d | n<br>d | n<br>d | n<br>d | + | + | + |
|  | *TPM1* | OG5_127228 | + | n<br>d | + | + | + | + | + | + | + | + | + |
| Plasmogamy | *CD9* | OG5_136376 | nd | n<br>d | n<br>d | n<br>d | n<br>d | n<br>d | n<br>d | n<br>d | n<br>d | n<br>d | n<br>d |
|  | *FIG1a* | OG5_138630 | nd | n<br>d | n<br>d | n<br>d | n<br>d | n<br>d | n<br>d | n<br>d | n<br>d | n<br>d | n<br>d |
|  | *FUS1a* | OG5_156961 | nd | n<br>d | n<br>d | n<br>d | n<br>d | n<br>d | n<br>d | n<br>d | n<br>d | n<br>d | n<br>d |
|  | *FUS2a* | OG5_171667_2168<br>47 | nd | n<br>d | n<br>d | n<br>d | n<br>d | n<br>d | n<br>d | n<br>d | n<br>d | n<br>d | n<br>d |
|  | *IZUMO1a* | OG5_151190 | nd | n<br>d | n<br>d | n<br>d | n<br>d | n<br>d | n<br>d | n<br>d | n<br>d | n<br>d | n<br>d |
|  | *KEX2* | OG5_127788 | nd | + | n<br>d | + | + | + | + | + | + | + | + |
|  | *MYO2* | OG5_126577 | + | + | + | + | + | + | + | + | + | + | + |
| Function | Gene | OG | *Arcella intermediata* | YT12-12 | YT9-10 | YT1-1 | YT5-6 | YT7-8 | YT4-5 | YT2-3 | YT8-9 | YT6-7 | YT10-11 |
|  | *PRM1a* | OG5_136560 | nd | n<br>d | n<br>d | n<br>d | n<br>d | n<br>d | n<br>d | n<br>d | n<br>d | n<br>d | n<br>d |
|  | *RVS161* | OG5_129890 | nd | n<br>d | n<br>d | n<br>d | n<br>d | n<br>d | n<br>d | n<br>d | n<br>d | n<br>d | n<br>d |
| Nuclear congression | *BIK1a* | OG5_139423 | nd | n<br>d | n<br>d | n<br>d | n<br>d | n<br>d | n<br>d | n<br>d | n<br>d | n<br>d | n<br>d |
|  | *CDC4* | OG5_129129 | + | n<br>d | + | + | n<br>d | + | n<br>d | + | + | + | + |
|  | *CDC28* | OG5_126712 | + | + | + | + | + | + | + | + | + | + | + |
|  | *CDC34* | OG5_129084 | + | n<br>d | n<br>d | + | + | + | n<br>d | + | + | + | + |
|  | *CDC37a* | OG5_133705 | nd | n<br>d | n<br>d | n<br>d | n<br>d | n<br>d | n<br>d | n<br>d | n<br>d | n<br>d | n<br>d |
|  | *CIK1a* | OG5_147436 | nd | n<br>d | n<br>d | n<br>d | n<br>d | n<br>d | n<br>d | n<br>d | n<br>d | n<br>d | n<br>d |
|  | *CIN1a* | OG5_156139 | nd | n<br>d | n<br>d | n<br>d | n<br>d | n<br>d | n<br>d | n<br>d | n<br>d | n<br>d | n<br>d |
|  | *CIN2a* | OG5_132672 | nd | n<br>d | n<br>d | n<br>d | n<br>d | n<br>d | n<br>d | n<br>d | n<br>d | n<br>d | + |
|  | *CIN4* | OG5_128319 | + | n<br>d | n<br>d | + | + | + | + | + | + | + | + |
|  | *KAR3* | OG5_138757 | nd | n<br>d | n<br>d | n<br>d | n<br>d | n<br>d | n<br>d | n<br>d | n<br>d | n<br>d | n<br>d |
| Function | Gene | OG | *Arcella intermedi*<br>*a* | YT12-12 | YT9-10 | YT1-1 | YT5-6 | YT7-8 | YT4-5 | YT2-3 | YT8-9 | YT6-7 | YT10-11 |
|  | *KAR4* | OG5_130096 | + | n<br>d | n<br>d | n<br>d | n<br>d | n<br>d | n<br>d | n<br>d | + | + | + |
|  | *KAR9a* | OG5_156957 | nd | n<br>d | n<br>d | n<br>d | n<br>d | n<br>d | n<br>d | n<br>d | n<br>d | n<br>d | n<br>d |
|  | *KEM1* | OG5_126774 | + | n<br>d | n<br>d | n<br>d | n<br>d | n<br>d | n<br>d | n<br>d | + | + | + |
| Karyogamy | *JEM1a* | OG5_163504 | nd | n<br>d | n<br>d | n<br>d | n<br>d | n<br>d | + | + | n<br>d | n<br>d | n<br>d |
|  | *KAR2* | OG5_126588 | + | + | + | + | + | + | + | + | + | + | + |
|  | *SEC63* | OG5_128413 | + | n<br>d | + | + | n<br>d | n<br>d | + | n<br>d | + | + | + |
|  | *SEC66a* | OG5_135424 | nd | n<br>d | n<br>d | n<br>d | n<br>d | n<br>d | n<br>d | n<br>d | n<br>d | n<br>d | n<br>d |
|  | *SEC72a* | OG5_136566 | nd | n<br>d | n<br>d | n<br>d | n<br>d | n<br>d | n<br>d | n<br>d | n<br>d | n<br>d | n<br>d |
| Bouquet formation | *SAD1* | OG5_129856 | nd | n<br>d | n<br>d | n<br>d | n<br>d | n<br>d | n<br>d | n<br>d | n<br>d | n<br>d | n<br>d |
| Cell cycle regulation | *CDC20* | OG5_126765 | + | n<br>d | + | n<br>d | n<br>d | + | n<br>d | + | + | + | + |
| DNA damage sensing/response | *MEC1/ATR* | OG5_128386 | + | n<br>d | n<br>d | n<br>d | n<br>d | n<br>d | n<br>d | + | + | + | + |
|  | *MRE11* | OG5_127969 | + | n<br>d | n<br>d | n<br>d | n<br>d | n<br>d | n<br>d | n<br>d | + | + | + |

| Function | Gene | OG | *Arcella*<br>*intermedia* | Meiosis-Reduced State<br>(MRS) |  |  |  |  | Meiosis-Enriched State<br>(MES) |  |  |  |  |
| --- | --- | --- | --- | --- | --- | --- | --- | --- | --- | --- | --- | --- | --- |
|  |  |  |  | YT12<br>-12 | YT9-10 | YT1-1 | YT5-6 | YT7-8 | YT4-5 | YT2-3 | YT8-9 | YT6-7 | YT10<br>-11 |
|  | *RAD1/MEI9* | OG5_130438 | nd | n<br>d | n<br>d | n<br>d | n<br>d | n<br>d | n<br>d | n<br>d | + | + | + |
|  | *RAD17* | OG5_129387 | nd | n<br>d | n<br>d | n<br>d | n<br>d | n<br>d | n<br>d | n<br>d | + | n<br>d | + |
|  | *RAD23* | OG5_127320 | + | n<br>d | n<br>d | n<br>d | n<br>d | n<br>d | n<br>d | + | + | + | + |
|  | *RAD24* | OG5_126706 | + | + | n<br>d | + | + | + | + | + | + | + | + |
|  | *RAD50* | OG5_127792 | + | n<br>d | n<br>d | n<br>d | n<br>d | n<br>d | n<br>d | + | + | + | + |
|  | *TEL1/ATM* | OG5_128955 | + | n<br>d | n<br>d | n<br>d | n<br>d | n<br>d | n<br>d | + | + | + | + |
|  | *XRS2* | OG5_180328 | nd | n<br>d | n<br>d | n<br>d | n<br>d | n<br>d | n<br>d | n<br>d | n<br>d | n<br>d | n<br>d |
| Double-strand<br>break formation | *REC114* | OG5_142109 | nd | n<br>d | n<br>d | n<br>d | n<br>d | n<br>d | n<br>d | n<br>d | n<br>d | n<br>d | n<br>d |
| Double-strand<br>break repair | *KU70* | OG5_129086 | + | n<br>d | n<br>d | n<br>d | n<br>d | n<br>d | n<br>d | + | + | + | + |
|  | *KU80* | OG5_129372 | + | n<br>d | n<br>d | n<br>d | n<br>d | n<br>d | n<br>d | n<br>d | + | + | + |
|  | *LIG4/DNL1* | OG5_130132 | + | n<br>d | n<br>d | n<br>d | n<br>d | n<br>d | n<br>d | n<br>d | + | + | + |
|  | *XRCC4/LIF1* | OG5_135131 | nd | n<br>d | n<br>d | n<br>d | n<br>d | n<br>d | n<br>d | n<br>d | n<br>d | + | n<br>d |

|  |  |  |  | Meiosis-Reduced State (MRS) |  |  |  |  | Meiosis-Enriched State (MES) |  |  |  |  |
| --- | --- | --- | --- | --- | --- | --- | --- | --- | --- | --- | --- | --- | --- |
| Function | Gene | OG | *Arcella intermediata* | YT12-12 | YT9-10 | YT1-1 | YT5-6 | YT7-8 | YT4-5 | YT2-3 | YT8-9 | YT6-7 | YT10-11 |
| *Recombinational repair* | *BRCA1* | OG5_159932 | nd | n<br>d | n<br>d | n<br>d | n<br>d | n<br>d | n<br>d | n<br>d | n<br>d | n<br>d | n<br>d |
|  | *BRCA2* | OG5_131863 | + | n<br>d | n<br>d | n<br>d | n<br>d | n<br>d | n<br>d | n<br>d | n<br>d | n<br>d | n<br>d |
|  | *DNA2* | OG5_129631 | + | n<br>d | n<br>d | n<br>d | n<br>d | + | n<br>d | n<br>d | + | + | + |
|  | *EXO1* | OG5_127511 | + | n<br>d | n<br>d | n<br>d | n<br>d | n<br>d | n<br>d | + | + | + | + |
|  | *FEN1* | OG5_127472 | + | n<br>d | n<br>d | n<br>d | n<br>d | n<br>d | n<br>d | + | + | + | + |
|  | *MLH1* | OG5_127201 | + | + | + | + | + | + | + | + | + | + | + |
|  | *MLH2* | OG5_202562 | nd | n<br>d | n<br>d | n<br>d | n<br>d | n<br>d | n<br>d | n<br>d | n<br>d | n<br>d | n<br>d |
|  | *MLH3* | OG5_130552 | + | + | + | + | + | + | + | + | + | + | + |
|  | *MMS4/EME1* | OG5_135664 | nd | n<br>d | n<br>d | n<br>d | n<br>d | n<br>d | n<br>d | n<br>d | + | + | + |
|  | *MPH1/FANCM* | OG5_128649 | + | n<br>d | n<br>d | n<br>d | n<br>d | n<br>d | n<br>d | n<br>d | + | n<br>d | + |
|  | *MSH2* | OG5_127538 | + | + | + | + | + | + | + | + | + | + | + |
|  | *MSH3* | OG5_130351 | + | + | + | + | + | + | + | + | + | + | + |
|  | *MSH6* | OG5_126895 | + | + | + | + | + | + | + | + | + | + | + |

| Function | Gene | OG | *Arcella*<br>*intermedia* | Meiosis-Reduced State<br>(MRS) |  |  |  |  | Meiosis-Enriched State<br>(MES) |  |  |  |  |
| --- | --- | --- | --- | --- | --- | --- | --- | --- | --- | --- | --- | --- | --- |
|  |  |  |  | YT12<br>-12 | YT9-10 | YT1-1 | YT5-6 | YT7-8 | YT4-5 | YT2-3 | YT8-9 | YT6-7 | YT10<br>-11 |
|  | *MUS81* | OG5_129162 | nd | n<br>d | n<br>d | n<br>d | n<br>d | n<br>d | n<br>d | + | + | + | + |
|  | *PMS1* | OG5_128001 | + | + | + | + | + | + | + | + | + | + | + |
|  | *PMS2* | OG5_128001 | + | + | + | + | + | + | + | + | + | + | + |
|  | *RAD51* | OG5_126834 | + | n<br>d | n<br>d | n<br>d | n<br>d | n<br>d | n<br>d | + | + | + | + |
|  | *RAD52* | OG5_130806 | + | n<br>d | n<br>d | n<br>d | n<br>d | n<br>d | n<br>d | n<br>d | n<br>d | + | n<br>d |
|  | *RAD54* | OG5_127098 | + | + | + | + | + | + | + | + | + | + | + |
|  | *RTEL1* | OG5_127294 | + | + | + | + | + | + | + | + | + | + | + |
|  | *SAE2* | OG5_138817 | nd | n<br>d | n<br>d | n<br>d | n<br>d | n<br>d | n<br>d | n<br>d | n<br>d | n<br>d | n<br>d |
|  | *SGS1* | OG5_126644 | + | + | + | + | + | + | + | + | + | + | + |
|  | *SLX1* | OG5_128732 | + | n<br>d | n<br>d | n<br>d | n<br>d | n<br>d | n<br>d | + | + | + | + |
|  | *SLX4* | OG5_136855 | nd | n<br>d | n<br>d | n<br>d | n<br>d | n<br>d | n<br>d | n<br>d | n<br>d | n<br>d | n<br>d |
|  | *SMC5* | OG5_128615 | + | n<br>d | n<br>d | n<br>d | n<br>d | n<br>d | n<br>d | n<br>d | + | + | + |
|  | *SMC6* | OG5_127751 | + | n<br>d | n<br>d | n<br>d | n<br>d | n<br>d | n<br>d | n<br>d | + | + | + |

|  |  |  |  | Meiosis-Reduced State (MRS) |  |  |  |  | Meiosis-Enriched State (MES) |  |  |  |  |
| --- | --- | --- | --- | --- | --- | --- | --- | --- | --- | --- | --- | --- | --- |
| Function | Gene | OG | *Arcella intermedia* | YT12-12 | YT9-10 | YT1-1 | YT5-6 | YT7-8 | YT4-5 | YT2-3 | YT8-9 | YT6-7 | YT10-11 |
|  | *YEN1* | OG5_132593 | nd | n<br>d | n<br>d | n<br>d | n<br>d | n<br>d | n<br>d | n<br>d | n<br>d | n<br>d | n<br>d |
| *Sister chromatid cohesion* | *PDS5* | OG5_128901 | + | n<br>d | n<br>d | n<br>d | n<br>d | n<br>d | n<br>d | n<br>d | n<br>d | + | + |
|  | *RAD21* | OG5_129513 | + | n<br>d | n<br>d | n<br>d | n<br>d | n<br>d | n<br>d | n<br>d | n<br>d | + | + |
|  | *SCC3* | OG5_127983 | + | n<br>d | n<br>d | n<br>d | n<br>d | n<br>d | n<br>d | n<br>d | n<br>d | + | n<br>d |
|  | *SMC1* | OG5_127449 | + | + | + | + | + | + | + | + | + | + | + |
|  | *SMC2* | OG5_127360 | + | + | + | + | + | + | + | + | + | + | + |
|  | *SMC3* | OG5_127789 | + | + | + | + | + | + | + | + | + | + | + |
|  | *SMC4* | OG5_127440 | + | + | + | + | + | + | + | + | + | + | + |
| **Total detected** |  |  | **59** | **24** | **26** | **28** | **28** | **29** | **29** | **43** | **58** | **61** | **61** |

A plus sign (+) indicates the gene was detected; nd indicates it was not detected. Gene names, OG identifiers, and functional categories follow the curated orthogroup mapping; presence/absence calls are taken from the BlastP results. Ref. shows detection in the reference *Arcella intermedia*. Isolates are sorted by total genes detected and grouped into Meiosis-Reduced (MRS) and Meiosis-Enriched (MES) states.

